# Neural Changes Underlying Rapid Fly Song Evolution

**DOI:** 10.1101/238147

**Authors:** Yun Ding, Joshua L. Lillvis, Jessica Cande, Gordon J. Berman, Benjamin J. Arthur, Min Xu, Barry J. Dickson, David L. Stern

**Affiliations:** Janelia Research Campus, Howard Hughes Medical Institute, 19700 Helix Drive, Ashburn, VA 20147, USA; Department of Biology, Emory University, Atlanta, GA 30322. USA

## Abstract

The neural basis for behavioural evolution is poorly understood. Functional comparisons of homologous neurons may reveal how neural circuitry contributes to behavioural evolution, but homologous neurons cannot be identified and manipulated in most taxa. Here, we compare the function of homologous courtship song neurons by exporting neurogenetic reagents that label identified neurons in *Drosophila melanogaster* to *D. yakuba*. We found a conserved role for a cluster of brain neurons that establish a persistent courtship state. In contrast, a descending neuron with conserved electrophysiological properties drives different song types in each species. Our results suggest that song evolved, in part, due to changes in the neural circuitry downstream of this descending neuron. This experimental approach can be generalized to other neural circuits and therefore provides an experimental framework for studying how the nervous system has evolved to generate behavioural diversity.

## Main text

Closely related animal species exhibit diverse behaviours, indicating that the nervous system can evolve rapidly to generate new adaptive behaviours. Little is known about the neuronal mechanisms underlying behaviour evolution. The fundamental organization of brains and neural circuits are largely conserved between related animals, suggesting that new behaviours may evolve mostly through modifications of existing neural circuitry^1^. Functional comparisons of homologous neurons may therefore illustrate how neural circuits evolve to cause behavioural diversification^2^.

We have explored this problem in species closely related to *Drosophila melanogaster*, allowing study of neural function underlying diverse behaviours within a taxon with a well-defined phylogenetic history^3^. Recently, neurons underlying many behaviours have been identified in *D. melanogaster*^4^, mainly through progress in targeting genetic reagents to small subsets of cell types^5^. The amenability of other *Drosophila* species to genetic manipulation^6^ provides a rare opportunity to perform functional comparisons of homologous circuits. In this study, we introduced neurogenetic reagents into non-*melanogaster Drosophila* species to study the function of homologous neurons in species that produce divergent courtship songs.

### Courtship song evolution in the *D. melanogaster* species subgroup

*Drosophila* species display sophisticated courtship rituals often involving a male chasing the female, dancing around her, singing to her by vibrating one or both wings, waving their sometimes-spotted wings, licking the female, and other behaviours^7^. Here we focus on singing, which can be systematically quantified more easily than most courtship behaviours^8^. Females detect courtship song through vibrations of their antennal arista and they mate preferentially with males that sing intact^9^ courtship song of their own species^10^. Courtship songs evolve rapidly, presumably as a result of female-choice sexual selection, and every species sings a unique song^7^.

*D. melanogaster* courtship song contains two basic elements: trains of pulses (pulse song) and continuous hums (sine song)^10^. Males of *D. yakuba* and *D. santomea*, however, do not sing sine song, but they produce two distinct modes of pulse song: thud song and clack song^11^. Thud song is generated by unilateral wing vibration; while clack song is generated when males vibrate both wings behind them (Fig. 1a). To infer the evolutionary origins of these song types, we surveyed song in all *D. melanogaster* subgroup species by simultaneously recording acoustic signals and fly movements during courtship. As reported previously^12,13^, all of these species except *D. orena* produce a pulse-like song by unilateral wing vibration (Fig. 1b, Extended Data Fig. 1, and SI Movie 1), suggesting that unilateral pulse song was produced by the common ancestor of the group. We therefore reclassified thud song as pulse song, because it is similar to the ancestral unilateral pulse type. Clack song appears to be an evolutionarily new or elaborated song that evolved in the common ancestor of *D. yakuba* and *D. santomea*.

**Figure 1:**
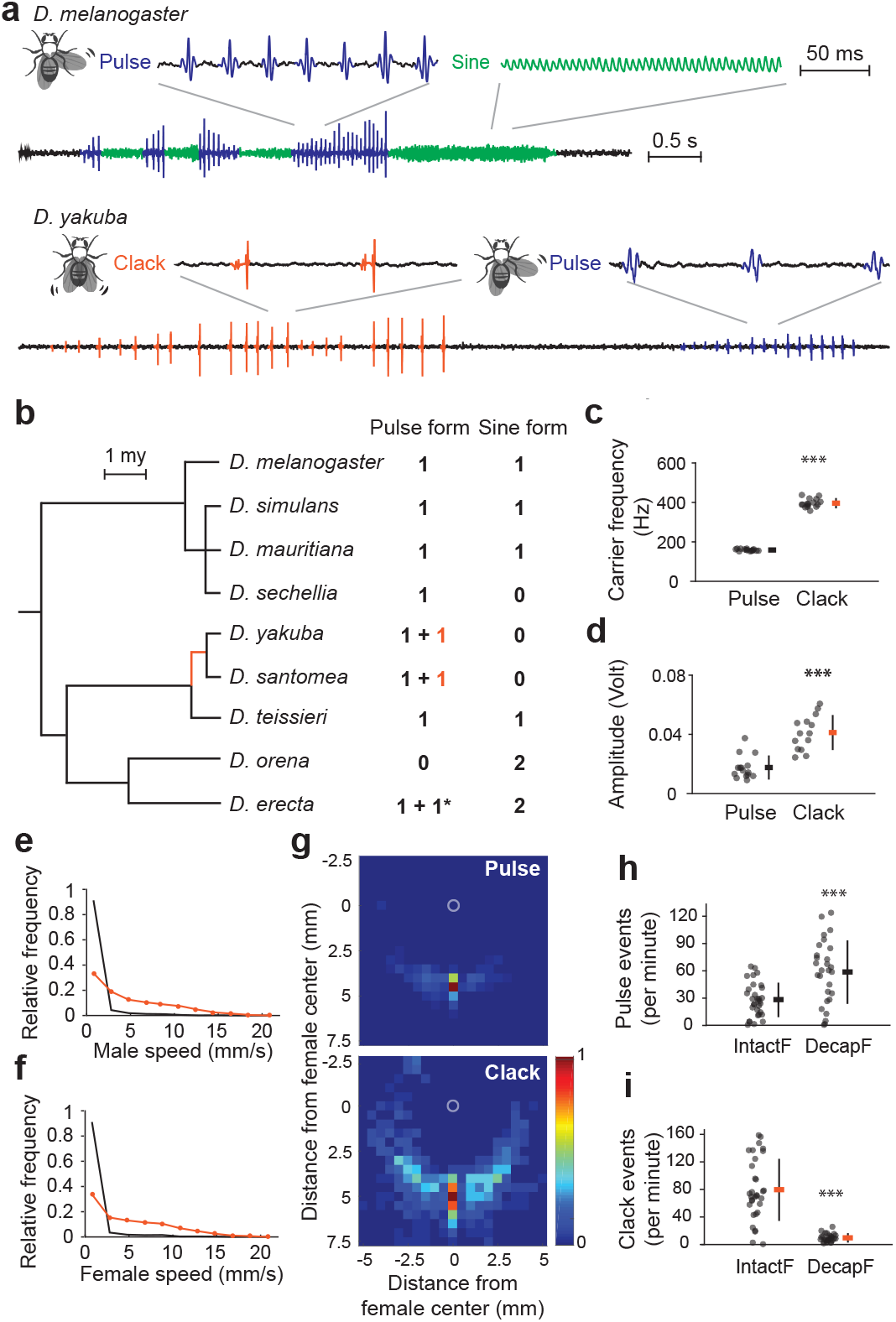
Divergence of courtship song between *D. melanogater* and *D. yakuba*. **a**, Illustration of courtship song of *D. melanogaster* and *D. yakuba*. Pulse and sine song are both generated via unilateral wing extensions, and clack song is generated by double wing vibrations. **b**, Evolution of primary male courtship song types in the *D. melanogaster* species subgroup. Almost all species produce pulse song via unilateral wing vibration. *D. yakuba* and *D. santomea* produces clack song, a distinct pulse song form by vibrating both wings behind the body (orange). *D. erecta* produces an additional song type composed of polycyclic pulses by vibrating one, occasionally two, wing(s) behind the body (marked by asterisk). The details of song types are shown in Extended Data Fig. 1. **c, d**, Carrier frequency (**c**) and amplitude (**d**) of pulse and clack song in *D. yakuba*. **e, f**, Probability density of male (**e**) and female speed (**f**) when the males are singing pulse (black) and clack (orange) song. n = 6. **g**, A heat map showing the position of male centroids relative to a centered female centroid (0,0) at the time of pulse song (top, n = 5, events = 1202) versus clack song (bottom, n = 5, events = 1222) in *D. yakuba*. Color represents the relative density of males located within a 0.25 mm^2^ square unit and the map is normalized relative to the square with highest density. **h, i**, Number of pulse (h) and clack (i) events across the tested recording session using intact (IntactF) and decapitated (DecapF) females. For panels **c, d, h**, and **i**, data for each animal and mean ± SD are shown. n > 16. *P* values estimated for one-way ANOVA using a permutation test. ***, *P* < 0.001.

*D. yakuba* clack song has a higher carrier frequency (Fig. 1c) and is louder (Fig. 1d) than pulse song^11,14^. Males sing clack song when both the male and female are moving faster than when males sing pulse song (Fig. 1e, f). Additionally, males sing pulse song mostly when they are located directly behind females, whilst they sing clack song across a wide range of distances and positions relative to females (Fig. 1g)^11^. Consistent with these observations, removing motion signals by providing males with a motionless decapitated female eliminated clack song but not pulse song (Fig. 1h, i). Thus, clack song is a high frequency, high amplitude song generated often during chasing. Pulse song in *D. yakuba*, by contrast, is quieter and generated when females slow down and allow males to follow them closely.

### Neurogenetic reagents from *D. melanogaster* label homologous neurons in *D. yakuba*

Several neurons required for *D. melanogaster* song have been identified^4^. P1 is a cluster of approximately 20 male-specific neurons per brain hemisphere that integrate multiple sensory stimuli^15–17^ and whose transient activity triggers a persistent courtship state^18,19^. Artificial activation of P1 neurons thus causes isolated males to produce many courtship behaviours, including song^15,16^. pIP10 is a single male-specific descending neuron per hemisphere that projects from the brain to the ventral nervous system (VNS) where it arborizes within the wing neuropil. In *D. melanogaster*, pIP10 acts as a command-like pathway to drive pulse song^15^.

We first examined whether the homologous neurons of P1 and pIP10 could be identified in non*-melanogaster* fly species. Here, we considered three criteria to define neurons as homologs. Homologous neurons (1) should be anatomically similar, (2) should express genetic markers that reflect a similar developmental origin, and (3) may be required to produce similar behaviours.

We first tested a subset of *D. melanogaster* GAL4 reagents that express in P1 and pIP10 neurons by integrating them into defined landing sites in *D. yakuba^6^*, and found that these GAL4 lines usually drove similar global expression patterns in both species (Extended Data Fig. 2). Since GAL4 reagents often drive expression in many unrelated neurons, we adopted the split-GAL4 strategy^5^ to identify reagents that labeled targeted neurons more cleanly. Because both the P1 and pIP10 neurons express the male-specific isoform of the sex-determination transcription factor-encoding gene *fruitless* (*fru*)^15^, we also generated *fru* expressing reagents in *D. yakuba* by replacing the first exon of the male-specific *fru* isoform with GAL4, GAL4 activating domain (AD), and DNA-binding domain (DBD) via CRISPR/Cas9-mediated homology dependent repair (HDR)^20^ (Extended Data Fig. 3a-c).

We screened two large *D. melanogaster* GAL4 driver line collections^21,22^ and identified split-GAL4 combinations that labeled P1 (GMR071G01-AD n VT054805-DBD, VT059450-AD n VT054805-DBD) and pIP10 (VT040556-AD n VT043047-DBD) with little extraneous expression (Extended Data Fig. 3d, f). In *D. yakuba*, we tested five and seven split-GAL4 combinations for P1 and pIP10 respectively. In all cases, we identified neurons with projection patterns similar to the targeted neurons (Extended Data Fig. 3e, g). We used multiple relatively clean P1 and pIP10 reagents for further behavioural analysis. Among them, the P1 reagent R71G01-AD n R15A01-DBD and the pIP10 reagent VT040346-AD n VT040556-DBD labeled male-specific neurons with the expected projection patterns almost exclusively. We show below that these labeled neurons also participate in producing courtship song in *D. yakuba*. Thus, based on criteria of anatomical similarity, expression of the same genetic markers (inferred because the male-specific neurons are labeled with the same GAL4 lines), and behavioural phenotypes, the labeled *D. yakuba* neurons appear to represent homologs of P1 and pIP10. We exploited these reagents to explore the circuitry changes contributing to song evolution.

### P1 neurons drive a persistent courtship state in both *D. melanogaster* and *D. yakuba*

We first tested whether P1 neurons have a conserved role in the two species. We expressed the red-shifted channel rhodopsin CsChrimson^23^ in *D. melanogaster* and *D. yakuba* P1 neurons and exposed isolated males to red light. Consistent with previous reports^15–19^, we found that optogenetic activation of P1 neurons in *D. melanogaster* triggered multiple courtship behaviours, including both pulse and sine song (Extended Data Fig. 4a). Transient optogenetic activation of P1 neurons in *D. yakuba* caused isolated males to perform extended bouts of courtship behaviour, including abdomen quivering, wing rowing, wing scissoring, and song that consisted mainly of clacks (SI Movie 2 and Extended Data Fig. 4b). Since *D. yakuba* males produce pulse song mostly when they move close to females, and a moving object triggers P1-activated *D. melanogaster* males to court vigorously^17,24^, we provided optogenetically activated *D. yakuba* males with a recently anesthetized male *D. yakuba* and found that P1-activated males then produced large quantities of pulse song (Extended Data Fig. 4b). Thus, P1 neurons have retained a conserved role in eliciting a persistent courtship state in both species. In addition, since activation of P1 neurons in *D. yakuba* males never caused production of sine song, the neural connections downstream of P1 likely evolved to cause loss of sine song.

### pIP10, the pulse song command neuron in *D. melanogaster*, is required for clack but not pulse song in *D. yakuba*

To address the role of neurons that specifically drive courtship song, we examined pIP10 function in both species. pIP10 inhibition in *D. melanogaster* was previously shown to reduce wing extension during courtship^15^. We found that pIP10 inhibition in *D. melanogaster* using our new split-GAL4 line caused almost complete elimination of pulse song and a small reduction in sine song produced during normal courtship (Fig. 2a). In contrast, pIP10 inhibition in *D. yakuba* eliminated clack song consistently across different split-GAL4 drivers and neuronal inhibitors (Fig. 2b and Extended Data Fig. 5a-c). In addition, in some treatments, pIP10 inhibition resulted in a quantitative reduction of pulse song (Extended Data Fig. 5a, c). Therefore, pIP10 is essential for pulse song production in *D. melanogaster* and for clack song production in *D. yakuba*. pIP10 also contributes to quantitative levels of sine song in *D. melanogaster* and pulse song in *D. yakuba*. It appears that pIP10 has switched its role in *D. yakuba*, from a descending neuron required primarily for pulse song to a neuron required primarily for clack song. These evolutionary changes likely occurred in the common ancestor of *D. yakuba* and *D. santomea*, because inhibiting pIP10 activity in *D. santomea*, using a non-sparse GAL4 line that labels pIP10, also blocked clack song but not pulse song during courtship (Extended Data Fig. 5d).

**Figure 2:**
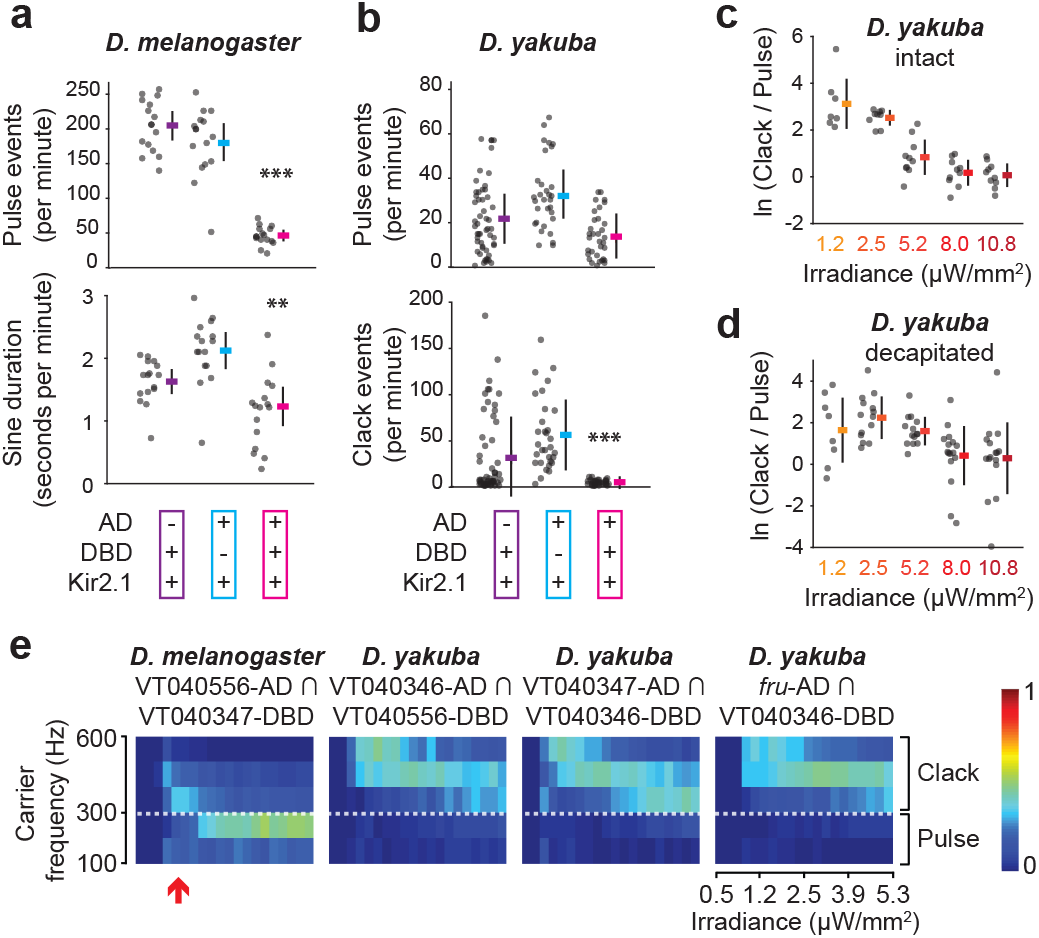
Different roles of pIP10 neurons in *D. melanogater* and *D. yakuba* courtship song. **a**, Pulse and sine song production of pIP10 silenced males (VT040556-AD ∩ VT040347-DBD > Kir2.1) in *D. melanogaster*. **b**, Pulse and clack song production of pIP10 silenced males (VT040346-AD ∩ VT040556-DBD > Kir2.1) in *D. yakuba. P* values estimated for one-way ANOVA using a permutation test. Significance is indicated only when the experimental group is significantly different from both control groups and the less significant result is shown. ***, *p* < 0.001; **, *p* < 0.01. n > 16. **c, d**, Natural log (ln) ratio of clack versus pulse events of pIP10 activated intact (c) and decapitated (**d**) males (VT040346-AD ∩ VT040556-DBD > CsChrimson) at different irradiances in *D. yakuba*. n > 8. Data for each animal and mean ± SD are shown. e, Heat map showing the distribution of song carrier frequency (pulse song for *D. melanogaster;* pulse and clack song combined for *D. yakuba)* of CsChrimson expressing pIP10 males with ramping irradiance using incremental steps of ~0.25 μW/mm^2^. Color represents the relative density of song events within a given carrier frequency range at the tested irradiance. For each genotype, mean of eight tested animals is shown. See Extended Data Fig. 6i for the results of each tested animal. In *D. yakuba*, pIP10 activation elicted mostly high frequency song events, presum-bly clack song, across all the tested irradiance levels. In *D. melanogaster*, pIP10 activation elicited high-frequency clack-like events (300-600 Hz) only at the very lowest irradiance levels (red arrow).

### Activation of pIP10 drives clack(-like) and pulse song in an intensity dependent manner in both species

To further explore the role of pIP10 in *D. yakuba*, we optogenetically activated pIP10 neurons expressing CsChrimson. In both intact and decapitated males, pIP10 activation drove clack and pulse song in an intensity dependent manner (Fig. 2c, d, Extended Data Fig. 6a, b, and SI Movie 3); low levels of pIP10 activation drove only clack song, but higher levels drove both clack and pulse song (Extended Data Fig. 6c, d). Sine song was never elicited. Artificially activated song had an inter-clack interval within the wildtype range (~115-140 ms)^11,14^ at low levels of light intensity, but an abnormally short inter-clack interval at higher activation levels (Extended Data Fig. 6e). Thus, the lower activation levels may more closely mimic natural pIP10 activity levels than the higher activation levels in *D. yakuba*.

The observation that pIP10 can drive both clack and pulse song in *D. yakuba* led us to re-examine the effect of activating pIP10 in *D. melanogaster*. Throughout most of the light intensity range, pIP10 activation drove mainly pulse song with normal carrier frequency (mode = ~200 Hz) (Extended Data Fig. 6f, g). Sine song was elicited rarely and the probability of sine song production increased with increasing light intensities (Extended Data Fig. 6h). Activation of pIP10 in *D. melanogaster* at very low light levels induced flies to produce a pulse song with a high carrier frequency (Extended Data Fig. 6f, g). We performed simultaneous recording of acoustic signals and fly movements and found that most, if not all, of these high frequency pulses are generated by double-wing vibrations, mimicking the wing posture of clack song (SI Movie 4). Similarly, optogenetic activation of P1 in *D. melanogaster* induced both normal pulses and pulses resembling clack song in carrier frequency (Extended Data Fig. 4c, d) and wing posture (SI Movie 5).

Overall, these observations are consistent with our inactivation experiments that revealed that pIP10 is required for clack song and natural levels of pulse song in *D. yakuba* and required for pulse song and natural levels of sine song in *D. melanogaster*. In addition, the induction of clack-like song by low levels of pIP10 stimulation in *D. melanogaster* suggests that the common ancestor of the *D. melanogaster* subgroup species possessed neural circuitry capable of producing clack song and that a circuit change in the common ancestor of *D. yakuba* and *D. santomea* allowed production of clack song more readily in these species. This evolutionary change was probably driven by female-choice sexual selection because *D. yakuba* males whose clack song is suppressed, by expression of Kir2.1 in pIP10 neurons, had significantly lower copulation success than control males (Extended Data Fig. 5e-g).

### Conserved physiological properties of pIP10 in *D. melanogaster* and *D. yakuba*

In both species, clack(-like) song is associated with low pIP10 activation levels. However, *D. melanogaster* produced clack-like song only in a narrow range of low activation levels, whereas *D. yakuba* produced clack song across a wide range of activation levels (Fig. 2e). Thus, the threshold between production of clack and pulse song has shifted between these species.

One hypothesis to explain these observations is that *D. melanogaster* pIP10 may exhibit higher excitability than *D. yakuba*, which would allow this neuron to more readily reach an activity level sufficient to drive pulse song. We therefore expressed CsChrimson in pIP10 neurons and assayed the responses of these neurons to light stimuli via ex vivo whole-cell patch-clamp recordings. Both species displayed increasing spiking rates with increasing levels of light stimulation, but we observed no statistically significant species-differences in spiking pattern at any illumination level (Fig. 3a, b) and the neurons responded very similarly in the illumination range corresponding to that used in the behavioural experiments (yellow range in Fig. 3b). We also found no differences in other electrophysiological properties, including responses to depolarizing current, resting membrane potential, spike threshold, spike amplitude, and afterhyperpolarization amplitude (Fig. 3c-i). Thus, while it is possible that there are species differences in electrophysiological properties that could not be measured at the soma, the results suggest that pIP10 electrophysiological properties are conserved between these species and cannot account for the differences in song type elicited by pIP10 activation. We also found that the onset of song in response to light stimuli is similar in both species (Fig. 2e) and red light penetrates the cuticle of both species similarly (Extended Data Fig. 7a). Thus, there are no technical differences in the ability to activate these neurons *in vivo* that can explain the species behavioural differences exhibited in response to pIP10 activation. These results suggest that the circuitry downstream of pIP10 has evolved to produce differential responses to similar pIP10 activity in these two species. However, we cannot exclude that species-specific differences in pIP10 neurotransmitter release, which we did not assay, may play a role in the behavioural differences elicited by similar levels of pIP10 activity.

**Figure 3:**
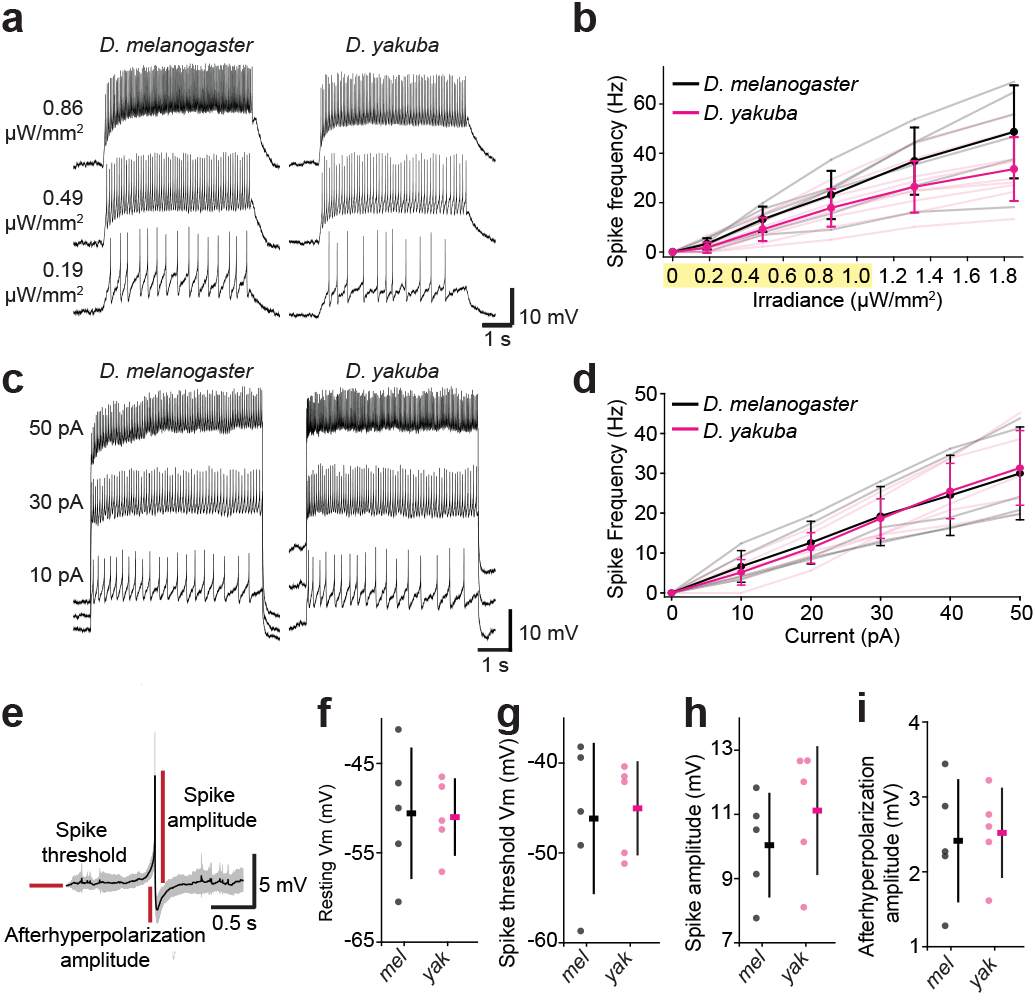
pIP10 has similar electrophysiology properties in ***D. melanogaster*** and *D. yakuba*. **a, b**, Spiking responses of CsChrimson expressing pIP10 neurons to different light irradiances in *D. melanogaster* and *D. yakuba*: representative examples of pIP10 spiking responses (**a**) and light dose-response curve of pIP10 (**b**). The irradiance range comparable to the range applied in the behaviour experiments is highlighted in yellow. **c, d**, Spiking responses of pIP10 to depolarizing current steps of different amplitudes in *D. melanogaster* and *D. yakuba:* representative examples of pIP10 spiking responses (**c**) and dose-response curve of pIP10 (**d**). For panel **a** and **c**, the resting membrane potential has been shifted to ease viewing. For panel **b** and **d**, data for each animal (translucent lines) and mean ± SD are shown. n = 5-9. Significance tested across species at each irradiance or current amplitude via two-way ANOVA and post-hoc Holm-Sidak pairwise multiple comparisons test. No significant differences. **e-h**, Comparision of pIP10 resting and spiking properties between *D. melanogaster (mel)* and *D. yakuba* (yak): resting membrane potential (**f**), spike threshold (**g**), spike amplitude (**h**), and afterhyperpolarization amplitude (**i**). Values for each animal and mean ± SD are shown. n = 5. Significance tested across species via t-test. No significant differences.

### Quantitative differences in pIP10 anatomy

One explanation for the species-specific difference in the song circuit response to pIP10 activity could be that pIP10 may synapse more extensively onto a conserved set of downstream neurons in *D. melanogaster* than in *D. yakuba*, thus resulting in an increased sensitivity to pIP10 activity in *D. melanogaster*. We do not yet know the synaptic partners of pIP10, so we cannot be sure that pIP10 connects with the same neurons in each species, but we can use the existing reagents to examine pIP10 arborization patterns to estimate the total number of synaptic connections in different regions.

Although the gross pIP10 projection patterns were similar in the two species (Fig. 4a, b, and SI Movie 6), we identified multiple species-specific differences (Fig. 4c-h). In the VNS, *D. yakuba* pIP10 displayed denser arbors than *D. melanogaster* pIP10 in several regions, particularly at the base of the mesothoracic triangle (red region 9 in Fig. 4f, h) and at the posterior-most descending projections in the T3 neuropil (blue region 12, magenta region 13 in Fig. 4f, h). Synaptotagmin staining, which marks pre-synaptic axons^25^, is observed in all of the pIP10 projections in the VNS^15^ (Fig. 4a and SI Movie 7), suggesting that pIP10 is largely, if not exclusively, presynaptic in the VNS. While the song circuit is incompletely known, all identified VNS song neurons co-localize with the mesothoracic triangle^9,15^. Thus, pIP10 pre-synaptic arbors are less dense in the region containing the song circuitry in *D. melanogaster* compared to *D. yakuba*, which suggests that total synaptic output alone cannot explain the species-specific effects of pIP10 activation. It is possible that the relative strength of pIP10 connectivity to downstream neurons has shifted between species, perhaps biasing the song circuit toward production of clack song in *D. yakuba*. Testing this hypothesis will require synaptic-level reconstruction of the song circuit in both species.

**Figure 4:**
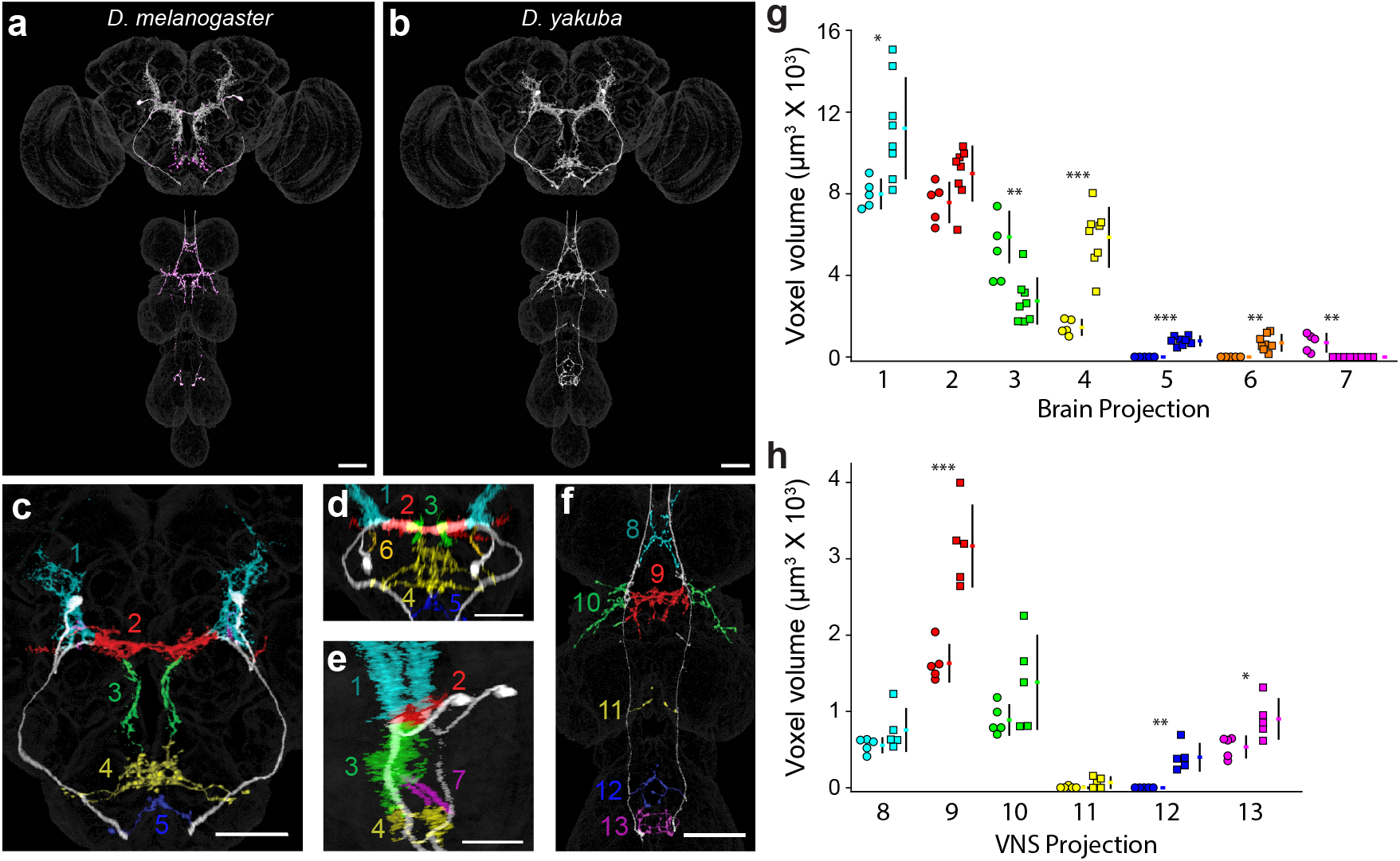
pIP10 anatomy in *D. melanogaster* and *D. yakuba*. **a, b**, Maximum intensity projections of representative segmented pIP10 neurons in both species (white) and synaptotagmin expression (magenta) in *D. melanogaster* aligned to a standard *D. melanogaster* brain and ventral nervous system (VNS). **c-f**, Brain (**c-e**) and VNS (**f**) arbor compartments that were further segmented for quantification represented in *D. yakuba* (**c, d, f**) or *D. melanogaster* (**e**). Quantified regions are labeled with separate colours and numbered. The lateral projection into the SEZ (**c**, blue region 5) and additional projections in the T3 neuropil of the VNS (**f**, blue region 12) are found only in *D. yakuba*. Shifted angle of the brain illustrates a projection near the soma found only in *D. yakuba* (**d**, orange region 6) and the posterior projection in the SEZ found only in *D. melanogaster* (**e**, magenta region 7). **g, h**, Quantification of arbor compartments for *D. melanogaster* (circles, left) and *D. yakuba* (squares, right). Colours and numbers in (**g**) and (**h**) correspond to numbers and coloured regions shown in (**c-e**) and (**f**), respectively. Scale bars represent 50 uM. *P* values measured via two-tailed t-tests for each arbor compartment. ***, p<0.001; **, p<0.01; *, p<0.05.

In addition to the differences found in the VNS, we also identified differences in pIP10 brain arborization patterns (Fig. 4). For example, *D. melanogaster* pIP10 extends arbors into the posterior portion of the SEZ that are not observed in *D. yakuba* (magenta region 7 in Fig. 4e, g). Conversely, *D. yakuba* displayed dense arbors extending laterally into the SEZ, whereas *D. melanogaster* displayed few projections in the same area^15^ (yellow region 4 in Fig. 4c-e, g). Synaptotagmin staining indicates that these SEZ arbors provide synaptic output in the brain (Fig. 4a and SI Movie 7). Thus, pIP10 anatomy has evolved in several ways in the brain as well, but further studies will be required to determine the evolutionary causes and functional consequences of these anatomical differences.

### Potential evolutionary antecedents of clack song in *D. melanogaster*

The observation that artificial activation of either P1 or pIP10 can, under certain conditions, elicit clack-like song in *D. melanogaster* males led us to re-examine the wildtype song of these species. Male *D. melanogaster* mainly produce low frequency (< 250 Hz) pulses, but they also sometimes produce high frequency (> 250 Hz) pulses (Extended Data Fig. 8a)^26^. Lower frequency pulses (150-250Hz) are associated with large wing extension angles (~60°-90°) and higher frequency pulses (250-500Hz) are sometimes associated with smaller wing extension angles (~20°-40°) and sometimes with larger wing extension angles^26^ (Extended Data Fig. 8d). We observed a negative correlation between wing angle and carrier frequency specifically for higher frequency pulses produced with shallow wing angles in *D. melanogaster* (Extended Data Fig. 8d) and this correlation is also observed in *D. simulans* and *D. mauritiana* (Extended Data Fig. 8e, f), suggesting that this reflects a conserved mechanism for generating high frequency pulses. We therefore searched more deeply for high frequency pulses generated by wings at acute angles and found, in ~1% of all pulse events, that *D. melanogaster* males generate short trains of high frequency pulses (mode = 298 Hz) without obvious extension of either wing (SI Movie 8). These song events are similar to the clack-like song elicited by P1 and pIP10 activation and suggest that the evolutionary antecedents of *D. yakuba* clack song can be observed rarely in other species. Thus, the evolution of abundant clack song in *D. yakuba* and *D. santomea* may reflect cooption of existing neural circuitry in the common ancestor of the *D. melanogaster* subgroup species through quantitative changes in VNS circuitry that increases the probability that pIP10 activity drives clack song.

## Discussion

Our results reveal five novel findings about the neural basis for the evolution of *Drosophila* courtship song. First, P1 neurons have retained a conserved function in establishing a persistent courtship state in *D. melanogaster* and *D. yakuba*. Since P1 activation results in species-specific songs, the sine song circuitry downstream of P1 has evolved between these species. Second, pIP10, a command neuron driving pulse song in *D. melanogaster* became largely dispensable for pulse song but essential for clack song in *D. yakuba*. Third, clack (-like) song and pulse song can be elicited by pIP10 activation in an intensity dependent manner in both species. This observation, together with our observation of rare *D. melanogaster* clack-like song, suggests that the common ancestor of these species possessed neural circuitry that could produce clack-like song. Fourth, differential responses to pIP10 activity, resulting in the production of mainly pulse song in *D. melanogaster* and clack song in *D. yakuba*, likely arose due to differences in neural circuitry downstream of pIP10. Finally, pIP10 neural anatomy has evolved both qualitatively and quantitatively, raising new questions about descending neuron evolution, structure and function.

It is curious that pIP10 drives mainly different songs in each species, when both species produce the apparently conserved pulse song. A similar observation has been reported in swim central pattern generator neurons of sea slugs, where homologous neurons play different roles in the production of homologous behaviours^27^. However, in both species pIP10 elicits primarily the louder song type in the context of males chasing females: pulse song in *D. melanogaster*^8^ and clack song in *D. yakuba* (Fig. 1d-f). In both species, when females slow down and allow males to follow them closely, they produce a quieter song, sine song in *D. melanogaster^8^* and pulse song in *D. yakuba* (Fig. 1d-f). Thus, the song types driven by increasing levels of pIP10 activity correlate with similar behavioural contexts in the two species (Fig. 5). This suggests that pIP10 receives similar inputs in both species that reflect information about the behavioural context and that downstream circuity has changed so as to elicit divergent songs that are appropriate to the social context.

**Figure 5:**
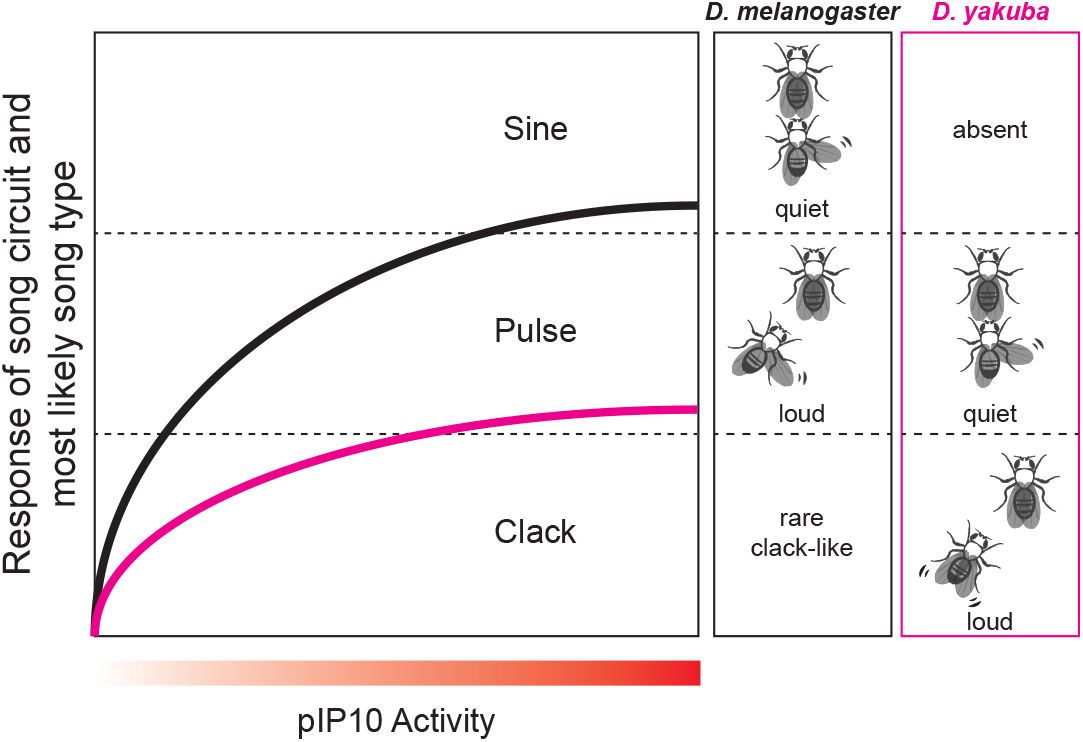
Evolving functional roles of pIP10 are associated with behavioral context, rather than specific song types. Increasing pIP10 activity drives a different distribution of songs in *D. melanogaster* and *D. yakuba*. In both species, low pIP10 activation drives loud songs and higher activation levels drive quiet songs. Very low levels of pIP10 activity can drive clack-like song in *D. melanogaster*, illustrating that this song type was likely present in the common ancestor, although abundant clack song is evolutionarily derived in *D. yakuba*. Evolution of alternative song types resulted mainly from changes in the song circuitry downstream of pIP10 activity.

In both species, pIP10 activation drives different song types in an intensity dependent manner. These observations are consistent with findings that minor differences in the activity of descending pathways can drive different patterns of song circuit activity in insects^29^. The availability of a descending pathway that can access different song circuit activity through minor modifications may have facilitated the rapid evolution of courtship behaviours in response to female choice sexual selection. In theory, evolution of the excitability of pIP10 could have driven evolution of new song types, but in this case, it has not. Instead, our data suggest that changes in the circuitry downstream of the descending inputs has evolved to generate the diverged patterns of courtship song in response to similar social cues.

*D. melanogaster* provides a powerful system for dissecting the neural circuitry underlying behaviour^30^ and the genus *Drosophila* contains over 1,500 species displaying divergent adaptive behaviours^31^. Our functional comparative approach illustrates how we can leverage tools developed in *D. melanogaster* to study the homologous neural circuitry underlying the evolution of many behaviours across this genus^32^.

## Acknowledgements

We thank Elizabeth Kim for extensive assistance in the laboratory, Kari Close, Christina Christoforou, and Gudrun Ihrke in the Janelia Project Technical Resources team for the assistance in dissection, histological preparation, and confocal imaging, Hideo Otsuna for assistance with brain registration and VVD Viewer, and Emily Robitschek and Jaime Cervantes for help in animal tracking using *ilastik*. Gerry Rubin and Heather Dionne for sharing plasmids, Artyom Kopp for providing the *D. orena* strain, and Mala Murthy for helpful comments on the manuscript.

## Author Contributions

Y.D., J.L.L., and D.L.S. designed the study and wrote the manuscript. Y.D. generated the *D. yakuba* and *D. santomea* genetic reagents with contributions from J.C. and D.L.S. J.L.L. and B.J.D. generated the *D. melanogaster* split-GAL4 lines. Y.D. and J.L.L. performed behavioural experiments. J.L.L. performed morphological quantification and electrophysiological experiments. B.A. built the behavioural rig and G.J.B. developed the k-means based song classification algorithm. M.X. performed fly husbandry and manual annotation of courtship. B.J.D. contributed to project discussion and manuscript editing.

## Supplementary Information

### Methods

#### Transgenic lines

In total, we generated 30 transgenic lines, including 7 UAS effector reagents, 12 GAL4 reagents, 8 split-GAL4 reagents, and 3 CRISPR/Cas9-mediated knock-in reagents in *D. yakuba* and *D. santomea*. Details are described in Supplementary Table 1. All transgenic injections were performed by Rainbow Transgenic Flies, *Inc* using standard protocols. Most of the plasmids were provided by Gerry Rubin.

#### Generation of *fru* GAL4, AD and DBD knock-in alleles

In general, the *fru* knock-in alleles were generated by precisely replacing the first exon of the male-specific *fru* isoform with GAL4, AD, and DBD fragment respectively via CRISPR/Cas9-mediated HDR. The GAL4, AD (p65ADZp), and DBD (ZpGAL4DBD) insertion fragments were amplified from the plasmids pBPGuW, pBPp65ADZpUw, and pBPZpGAL4DBDUw (Addgene), respectively. The donor plasmids were constructed by concatenating a 1.2 kb left homology arm, the insertion fragment (GAL4, AD or DBD), a 3XP3::DsRed marker (in inverted sequence), a 1.1 kb right arm, and a 1.8 kb backbone using Gibson Assembly^33^. A pair of guide RNAs (5’-GCGACGTCACAGGATTATTT-3’ and 5’-GGAGGCTTACCTAGGGGATG-3’) was cloned into the PCFD4 plasmid^34^. The donor plasmid, the PCFD4 plasmid, and *in vitro* transcribed *D. melanogaster* codon optimized Cas9 mRNA were co-injected into the embryos as described previously^35^.

#### Behavioural apparatus

The song recording apparatus was described previously^8^. A CMOS camera (Point Grey Flea3 1.3 MP Mono USB3) with F1.4 25mm lens (Navitar) was placed 22 cm above the behavioural chamber for video recording. Synchronization between audio and video was achieved by triggering each frame with a pulse generated by the same data acquisition board (National Instruments) that digitized the audio.

For CsChrimson stimulation, red LEDs (635 nm) were placed 6 cm above the behavioural chamber to provide the light stimulus. Pulse-width modulation with a 100 kHz frequency was used to adjust the light intensity with a custom-build high-side switching LED controller, a data acquisition board (National Instruments) was used to generate the timing signal for the light stimulus. The synchronization of this stimulus was achieved by recording the output of the controller on the same data acquisition board that digitized the audio. Infrared LEDs (850 nm) were used to provide illumination for video recording, and the lens was attached with an 800 nm long pass filter (Thorlabs) to remove light produced by the red LEDs.

The audio, video, and light stimulus data were captured using the custom Matlab program *omnivore* (<u>https://github.com/bjarthur/omnivore</u>). Data were visualized and movies were exported using the custom Matlab program *Tempo* (<u>https://github.com/JaneliaSciComp/tempo</u>).

#### Courtship song analysis

The *D. melanogaster* song was segmented using FlySongSegmenter^36^ with modified parameters^37^ to capture higher frequency pulse events. The *D. simulans* and *D. mauritiana* song was also segmented using FlySongSegmenter with different parameters^35^. For *D. yakuba* and *D. santomea* song, we developed an approach (https://github.com/gordonberman/FlySongClusterSegment) to automatically classify pulse and clack song. In brief, we sampled a subset of putative song events that are above noise threshold from a training set of *D. yakuba* and *D. santomea* wild-type songs, performed k-means clustering, and manually defined each cluster as template for pulse, clack, or noise. Models were constructed from each of these templates, such that a new event could be assigned a likelihood (p(event I template)). Data from subsequent songs were assigned to one of the song templates by finding the maximum likelihood template. Template assignments for the training set were manually checked to ensure accuracy. For each song type, we used a combination of “good” templates created from multiple training subsets and added additional templates until we were satisfied with the resulting assignments. All the *D. yakuba* and *D. santomea* songs were analyzed using the same set of templates. Courtship song elicited by P1 activation in *D. yakuba* was manually annotated, as large amount of other courtship behaviors elicited, including abdomen quivering and wing scissoring, generated sound that confounded automatic song identification. Clack-like song in wild-type *D. melanogaster* was manually annotated as train of pulses (n > 2) generated when males vibrated both wings behind the body.

Song parameters were measured using BatchSongAnalysis (<u>https://github.com/dstern/BatchSongAnalysis</u>) with modifications to analyze *D. yakuba* and *D. santomea* data. We characterized pulse carrier frequency in the following way. Individual pulse events represent acceleration of the wing from a stationary position to a maximum velocity and then deceleration of the wing to a stationary position again. Consequently, power of an individual pulse event can be distributed across a wide frequency range. We observed that the fast Fourier transform of individual pulses sometimes had a single strong peak at a single frequency and sometimes had power distributed across a range of frequencies. Thus, there does not appear to be an obviously single best way to measure the “carrier frequency” of a pulse event. We therefore measured pulse carrier frequency in two different ways and compared the results. First, we identified the dominant peak in the spectrogram. Second, we calculated the spectral centroid (“center of mass”) of the spectrogram, as recommended by Clemens et al^26^. The results from both methods produced qualitatively similar patterns and we show the results from analysis of the major peak of the fast Fourier transform in the figures.

#### Video analysis

All videos were recorded at a frame rate of 30 Hz. Automatic fly tracking was achieved using the pixel classification and animal tracking functions of *ilastik*^38^ and manually checked afterwards. The male and female speed during singing were measured as the distance of fly centers over a 200 ms time window centered by each song event. The relative positions of males to females were manually annotated by measuring the distance between their thoracic centers and the angle between their body axes. Frames were excluded from analysis if the male and female were positioned on opposite sides of the chamber or if one or both flies were positioned on a wall. Wing angle was manually annotated by measuring the angle between the thoracic center-wing distal end axis and the body axis. To establish the relationship between wing angle and pulse carrier frequency, we randomly measured the wing angle of one event among all the pulse events with the same frequency (closest integer) within the defined range *(D. melanogaster:* 150 - 500 Hz; *D. simulans* and *D. mauritiana:* 150 - 650 Hz), and four animals were scored for each species.

#### Behavioural experiments

To characterize song types of the *D. melanogaster* species subgroup (results shown in Fig. 1i and Extended Data Fig. 1), we used relatively larger chamber (2 cm X 4 cm) to capture a wider range of courtship dynamics than are normally observed in smaller chambers. A 4-10 day old virgin male (single housed) and a 3-5 day old virgin female (group housed) were placed into the chamber and recorded for 20-30 minutes. *D. sechellia, D. teissieri*, and *D. orena* males did not court much under this setting, so they were further recorded using a smaller round chamber (1 cm diameter) to collect more courtship events. For each species, At least 10 recordings with abundant song were collected. The strain information and the sample size for each species are described in Supplementary Table 2. All other song recordings in this study employed 1 cm diameter chambers.

For pIP10 inactivation in *D. melanogaster* and *D. yakuba*, experimental flies were obtained by crossing split-GAL4 males with females carrying a UAS line for a neuronal inhibitor (UAS-Kir2.1 or UAS-TNT), and control flies were obtained by crossing neuronal inhibitors to the corresponding AD or DBD lines alone. For the *D. melanogaster* split VT040556-AD n VT040347-DBD, males were crossed to Kir2.1 females (*w*+; UAS-Kir2.1). For the *D. yakuba* splits VT040346-AD; VT040556-DBD and VT040347-AD n VT040346-DBD, males were crossed with females carrying 3XP3::DsRed marked Kir2.1 in a wildtype background (yak*w*+; Kir2.1). This is not applicable for the split *fru*-AD; VT040346-DBD, because *fru*-AD is also marked with a 3XP3::DsRed and could not be maintained as homozygotes; therefore, males of this split were crossed to females carrying the neuronal inhibitor in a *white* mutant background (yak2180_Kir2.1 and yak2180_TNT). For pIP10 inactivation in *D. santomea*, experimental flies were obtained by crossing the GAL4 males (san2150_VT040556) with Kir2.1 females (san2174_Kir2.1), and crossing the GAL4 and Kir2.1 males with the *D. santomea* white females (with the same genetic background as the GAL4 and Kir2.1 flies) to generate the controls. For song recording, a 5-7 day old virgin male (single housed) and a < 1 day old virgin female (group housed) were placed into a 1 cm diameter chamber and recorded for 30 minutes. To measure copulation latency, a 4-7 day old virgin male (single housed) and a 4-7 day old virgin female (group housed) were placed in 1 cm diameter chambers and video recordings were collected for 60 minutes. Copulation time was scored manually. The control and experimental groups were always recorded simultaneously.

For CsChrimson experiments, the split-GAL4 males were crossed to females carrying UAS-CsChrimson *(D. melanogaster:* 20XCsChrimson-mVenus; *D. yakuba:* 20XCsChrimson-tdTomato). Males were collected 1-2 days after eclosion and group-housed in the dark for 6-7 days on standard media containing 0.5 mM trans-retinal (Sigma-Aldrich). Single isolated males were used for the experiments. To elicit pulse song for P1 activation in *D. yakuba*, we also included a 6-7 day old group-housed male to provide a moving object. Constant red light was applied for stimulation, and the specific protocols for each experiment are described in the following paragraph.

To activate P1 neurons, five replicates of a 30 s OFF and 30 s ON stimulation cycle were performed at the following light intensities from low to high: 2.5 μW/mm^2^, 5.3 μW/mm^2^, 8.0 μW/mm^2^, and 10.8 μW/mm^2^. A 5-minute resting period was provided before stimulation at the next intensity level. To activate pIP10 neurons, a stimulation cycle consists of 20 s OFF and 10 s ON period at the following light intensities from low to high: 1.2 μW/mm^2^, 2.5 μW/mm^2^, 5.3 μW/mm^2^, 8.0 μW/mm^2^, 10.8 μW/mm^2^, and 15.6 μW/mm^2^. This cycle was repeated ten times. The above two protocols were designed differently because the temporal dynamics of behavioural response to light stimulation differ between P1 and pIP10. P1 activation elicits courtship behaviour in a probabilistic manner: the elicited behaviour is not time locked to the stimulation and occurs with variable latencies^18,19^. We therefore allowed completion of behaviours in response to stimulation before increasing the intensity level. In contrast, pIP10 activation elicits courtship song acutely, so each cycle includes stimulation with a ramping intensity. For activating pIP10 neurons with a ramping intensity using small incremental step, a stimulation cycle consists of 10 s OFF and 5 s ON period at an intensity from 0.5 μW/mm^2^ to 5.3 μW/mm^2^ with an incremental step of ~ 0.25 μW/mm^2^. Four repeats of this cycle were performed.

During analysis of song phenotypes, outliers were systematically excluded in our song analysis pipeline using the Grubbs test with α=0.05 (<u>http://www.mathworks.com/matlabcentral/fileexchange/3961-deleteoutliers</u>). *P* values for ANOVAs were estimated with 10,000 permutations (<u>http://www.mathworks.com/matlabcentral/fileexchange/44307-randanova1</u>). For testing copulation latency, *P* values were calculated via a logrank test.

#### Immunostaining and imaging

The dissections, immunohistochemistry, and imaging of fly central nervous systems were done as described previously (Aso et al., 2014). In brief, brains and VNSs were dissected in Schneider’s insect medium and fixed in 2% paraformaldehyde (diluted in the same medium) at room temperature for 55 min. Tissues were washed in PBT (0.5% Triton X-100 in phosphate buffered saline) and blocked using 5% normal goat serum before incubation with antibodies. Tissues expressing GFP were stained with rabbit anti-GFP (ThermoFisher Scientific A-11122, 1:1000) or chicken anti-GFP (Abcam ab13970, 1:1200) and mouse anti-BRP hybridoma supernatant (nc82, Developmental Studies Hybridoma Bank, Univ. Iowa, 1:30), followed by Alexa Fluor® 488-conjugated goat antirabbit or goat anti-chicken and Alexa Fluor® 568-conjugated goat anti-mouse (ThermoFisher Scientific A-11034 and A-11031) or ATTO 647-conjugated goat antimouse (15048, Active Motif) antibodies, respectively. Tissues expressing tdTomato were stained with rabbit anti-DsRed (Clontech 632496, 1:1000) and nc82 (see above), followed by Cy^TM^3-conjugated goat anti-rabbit and Cy^TM^2-conjugated goat anti-mouse antibodies (Jackson ImmunoResearch 111-165-144 and 115-225-166), respectively.

For polarity staining, tissues expressing GFP and SYN::HA in pIP10, driven by the *D. melanogaster* split-GAL4 line VT040556-AD n VT040347-DBD, were stained with chicken anti-GFP (see above), rabbit anti-HA (Cell Signaling Technology #3724, 1:1000), and nc82 (see above), followed by goat anti-chicken AlexaFluor 488 conjugated (Thermo Fisher Scientific A-11039), Goat anti-rabbit, Cy5 conjugated (Jackson ImmunoResearch 111-175-144), and goat anti-mouse, AlexaFluor 568 conjugated (see above), respectively. After staining and post-fixation in 4% paraformaldehyde, tissues were mounted on poly-L-lysine-coated cover slips, cleared, and embedded in DPX as described. Image z-stacks were collected at 1 μm intervals using an LSM710 or LSM880 confocal microscope (Zeiss, Germany) fitted with a Plan-Apochromat 20x/0.8 M27 objective. Parent GAL4 images of *D. melanogaster* are from Jennet et al., 2012. Images of VT040346, VT040347, and VT040556 and all male split-GAL4, *D. yakuba*, and *D. santomea* images were generated by the FlyLight project team.

#### pIP10 segmentation and quantification

pIP10 split-GAL4 lines *(D. melanogaster:* VT040556-AD ∩ VT040347-DBD, *D. yakuba:* VT040436-AD ∩ VT040556-DBD) were registered using the Computational Morphometry Toolkit (<u>http://nitrc.org/projects/cmtk</u>)^39^ to the JFRC 2010 brain template and a newly generated VNS template. pIP10 neurons were segmented by extracting the pIP10 signal from non-target neuron expression using VVD Viewer (<u>https://github.com/takashi310/VVDViewer/blob/master/README.md</u>)^40,41^. Individual pIP10 arbors were then further segmented into unambiguous compartments to compare arbor volume across species (Fig. 4). Like arbors from the left and right pIP10 neurons were combined in each individual. In the brain, the medial arbors (red region 2) were defined as those extending from the medial, horizontal branch that is the most proximal projection to the soma, the dorsal arbors (cyan region 1) as those extending from the branch projecting dorsally from the medial branch, the ventral arbors (green region 3) as those extending from the branch projecting ventrally from the medial branch, the ventral-posterior arbors (magenta region 7, found only in *D. melanogaster*) as those extending from the branch projecting posteriorly from the ventral branch, the dorsal SEZ arbors (yellow region 4) as those extending from the two dorsal-most branches projecting from the descending projection into the SEZ, the ventral SEZ arbors (blue region 5, only found in *D. yakuba*) as those extending from the one ventral-most branch projecting from the descending projection into the SEZ, and the soma arbors (orange region 6, only found in *D. yakuba*) as those extending from the smaller, secondary branch proximal to the soma. In the VNS, the anterior triangle arbors (cyan region 8) were defined as those extending from the anterior most portion of the mesothoracic triangle projections, the medial triangle arbors (red region 9) as those medial to the descending projection at the base of the mesothoracic triangle projections, the lateral triangle arbors (green region 10) as those lateral to the descending projection at the base of the mesothoracic triangle projections, the T2 descending arbors (yellow region 11, only present in 1/5 *D. melanogaster* and 3/5 *D. yakuba*) as those extending from the descending projection in the T2 neuropil, and the T3 descending arbors (blue region 12 and magenta region 13) as those extending from the descending projection in the T3 neuropil. VVD Viewer was used to calculate the voxel volume of each of these compartments. Significance was determined via two-tailed t-tests of each compartment across species.

#### Cuticle light penetrance

An optical fiber (Thorlabs, FG105LVA) connected (S151C, Thorlabs) to a ThorLabs PM100D Compact Power and Energy Meter was positioned in front of a 3 mm LED fiber connected to a CoolLED pE4000 so that the LED would illuminate the front of the bare fiber (the only portion of the fiber that responded to light). Male flies were decapitated and the bare fiber was inserted into the head via the neck connective. 470, 525, 580, 595, and 635 nm constant on light pulses were presented in a quasi-random order. Light power for each wavelength was recorded once the measurement stabilized. The head was then carefully removed from the fiber and light was presented to the fiber again for measurement. Penetrance was calculated as the light power measurement in the head divided by the measurement without the head present. Measurements were similar to a previous *D. melanogaster* report^19^.

#### Irradiance calculation for behaviour and electrophysiology experiments

Irradiance was measured using a ThorLabs PM100D Compact Power and Energy Meter with a Console S130C Slim Photodiode Power Sensor. For behaviour experiments, the sensor (diameter, 9.5 mm) was positioned in the same location as the arena (diameter, 10.5 mm) directly over the recording chamber microphone. Irradiance was calculated as the raw light power measured divided by the area of the sensor (70.88 mm^2^).

For electrophysiology experiments, the 635 nm LED stimulus (pE4000, CoolLED) was delivered (with stacked 2.0 and 1.0 neutral density filters in the beam path) through a Zeiss Examiner Z1 with a W N-Achroplan 40X/0.75 water objective. The patched pIP10 soma was positioned in the center of the objective, and thus, in the center of the focused LED beam. The LED beam size was calculated using a beam profiler (WinCamD-UCD12, DataRay) with the sensor placed at approximately the same distance from the objective as the sample during experiments (2 mm). This yielded a 1/e^2^ beam area of 0.95 mm^2^. Light power was also measured with the sensor placed 2 mm away from the center of the objective. In an effort to measure the light power of the focused beam and reduce the amount of unfocused or reflected light from being measured by the 70.88 mm^2^ sensor, a painted black foil sheath was placed over the sensor with an opening for the objective to deliver light. Irradiance was calculated as the raw light power measured divided by the 0.95 mm^2^ focused beam area.

#### Electrophysiology

Male flies were collected shortly after eclosion and housed in isolation. 1-3 day old and 6-8 day old flies were tested. Because no age-related differences were found in any electrophysiological properties measured, flies within species were pooled. Individual flies were anesthetized by cooling. The brain and VNS were removed from the animal and placed into external saline composed of (in mM) 103 NaCl, 3 KCl, 5 N-tris(hydroxymethyl) methyl-2-aminoethane-sulfonic acid, 10 trehalose dihydrate, 10 glucose, 26 NaHCO_3_, 1 NaH_2_PO_4_, 4 MgCl_2_, 3 KCl, 2 sucrose, and 1.5 CaCl_2_ (280-290 mOsm, pH 7.3; components from Sigma Aldrich). The connective tissue and sheath were removed using fine forceps and the CNS was transferred to a chamber (Series 20 Chamber, Warner Instruments) superfused with external saline (carboxygenated with 95%O_2_ and 5%CO_2_) and held into place via a custom holder.

Fluorophore expressing pIP10 neurons were visualized using a Zeiss Examiner Z1 with a W N-Achroplan 40X/0.75 water objective, 470 nm or 580 nm LED illumination (pE-4000, CoolLED), and an IR-1000 infrared CCD monochrome video camera (Dage-MTI). The pIP10 soma was clearly identifiable as the only fluorophore expressing neuron in the region. Whole-cell recordings were obtained using glass patch electrodes filled with an internal solution composed of (in mM) 140 K-gluconate, 10 HEPES, 1 KCl, 4 MgATP, 0.5 Na_3_GTP, and 1 EGTA (270-280 mOsm, pH 7.3, components from Sigma Aldrich) connected to an Axopatch 700B amplifier (Molecular Devices) and digitized (10 kHz) with a Micro 1401-3 using Spike2 software (Cambridge Electronic Design). Glass electrodes were made using a P-1000 micropipette puller (Sutter) from borosilicate glass (Sutter; 1.2 mm outer diameter, 0.69 mm inner diameter). The pipette tip opening was less than one micron with a resistance between 5 and 15 MΩ.

pIP10 neurons were recorded in current clamp mode. The input resistance of pIP10 was tested intermittently throughout the recording and was above 600 MΩ for all data here. All data except for light dose-response recordings were obtained in pIP10 neurons expressing either 10XUAS-IVS-mCD8::GFP (*D. melanogaster*) or UAS-myr-GFP (*D. yakuba*). Light dose-response recordings were obtained in pIP10 neurons expressing 20XCsChrimson-mVenus (*D. melanogaster*) or 20XCsChrimson-tdTomato (*D. yakuba*). The spike threshold was determined as the lowest membrane potential at which pIP10 fired action potentials. Spike amplitude and afterhyperpolarization amplitude were measured for 5 spikes fired near threshold and averaged. pIP10 rested ~5 mV below spike threshold in both species and was held ~5 mV below spike threshold while light dose-response experiments were conducted. pIP10 was excited by a constant-on 5s depolarizing current steps or 635 nm light pulses delivered every 30 s through the objective while recording from the soma which was positioned in the center of the field of view. The light stimuli were similar to those used in behaviour experiments (both constant on stimuli; similar range of irradiance based on cuticle penetrance calculations). Light irradiance and current amplitude presentation order was varied from experiment to experiment. There was no indication that the order of intensity presentation affected the pIP10 response. Spikes were identified and counted using Spike2 scripts, and verified via manual inspection. Spike frequency (total spikes/5s) was plotted versus current step amplitude or light stimulus irradiance (SigmaPlot 12.5).

**SI Movie 1: Primary male courtship song types for all nine species in the *D. melanogaster* species subgroup**. For each song type in each species, 6-25 representative song clips were randomly chosen and concatenated for demonstration. The nomenclatures of song types are consistent with Extended Data Fig. 1.

**SI Movie 2: Courtship behaviors elicited by P1 CsChrimson activation in *D. yakuba*.** For each P1 split-GAL4 driver, representative examples of pulse song, clack song, abdomen quivering, wing rowing, and wing scissoring elicited by CsChrimson activation were shown.

**SI Movie 3: Courtship songs elicited by pIP10 CsChrimson activation with ramping irradiances in *D. yakuba*.** Expression of CsChrimson in pIP10 neuron was driven by the split-GAL4 line VT040346-AD ∩ VT040556-DBD. Light stimulation window and irradiance level were indicated in the video. Lower irradiance level elicited only clack song, featured by higher frequency pulses produced by vibration of both wings behind the body. Higher irradiance levels first elicited clack song and then pulse song, featured by lower frequency pulses produced by vibration of a single extended wing. The transition from clack to pulse occurred sooner with increasing irradiances.

**SI Movie 4: Clack-like song elicited by pIP10 CsChrimson activation with ramping irradiances in *D. melanogaster***. Expression of CsChrimson in pIP10 neuron was driven by the split-GAL4 line VT040556-AD ∩ VT040347-DBD. Light stimulation window and irradiance level were indicated in the video. The lower stimulation level mostly triggered clack-like song, and the higher stimulation level triggered pulse song with small amount of sine song.

**SI Movie 5: Clack-like song elicited by P1 CsChrimson activation in *D. melanogaster***. For each P1 split-GAL4 driver, two representative clips were randomly chosen and concatenated for demonstration.

**SI Movie 6: pIP10 anatomy in *D. melanogaster* and *D. yakuba***. Individual representative examples of pIP10 in *D. melanogaster* (green) and *D. yakuba* (magenta) registered and overlaid on a common *D. melanogaster* template brain and VNS.

**SI Movie 7: pIP10 synaptotagmin expression in *D. melanogaster***. Synaptotagmin expression in five registered *D. melanogaster* pIP10 neurons overlaid on a common *D. melanogaster* template brain and VNS. Each individual is a different color.

**SI Movie 8: Clack-like songs in wild-type *D. melanogaster***. Four representative song clips were randomly chosen and concatenated for demonstration.

**Supplementary Table 1:** Details of the transgenic lines generated in this study.

| Stock Name | Donor Plasmid | Fly Strain | Species | Method |
| --- | --- | --- | --- | --- |
| yakw; myrGFP | pBac{UAS-myrGFP, mw+} | UCSD 14021-0261.02 | *D. yakuba* | piggBac transgenesis |
| yakw+; Kir2.1 | pBac{UAS-EGFP::Kir2.1, 3XP3::DsRed} | UCSD 14021-0261.01 | *D. yakuba* | piggBac transgenesis |
| yak1730_CsChrimson | 20xUAS-CsChrimson::tdTomato | yak1730 | *D. yakuba* | attB/P integration |
| yak2180_Kir2.1 | pJFRC49 10XUAS-EGFP::Kir2.1 | yak2180 | *D. yakuba* | attB/P integration |
| yak2180_TNT | pJFRC34 5XUAS-TNT | yak2180 | *D. yakuba* | attB/P integration |
| san1504_CsChrimson | 20xUAS-CsChrimson::tdTomato | san1504 | *D. santomea* | attB/P integration |
| san2174_Kir2.1 | pJFRC49 10XUAS-EGFP::Kir2.1 | san2174 | *D. santomea* | attB/P integration |
| yak2180_R22D03 | R22D03-GAL4 | yak2180 | *D. yakuba* | attB/P integration |
| yak2177_R22D03 | R22D03-GAL4 | yak2177 | *D. yakuba* | attB/P integration |
| yak2180_R71G01 | R71G01-GAL4 | yak2180 | *D. yakuba* | attB/P integration |
| yak2177_R71G01 | R71G01-GAL4 | yak2177 | *D. yakuba* | attB/P integration |
| yak1730_R15A01 | R15A01-GAL4 | yak1730 | *D. yakuba* | attB/P integration |
| yak1664_R15A01 | R15A01-GAL4 | yak1664 | *D. yakuba* | attB/P integration |
| yak1664_VT040346 | VT040346-GAL4 | yak1664 | *D. yakuba* | attB/P integration |
| yak1694_VT040346 | VT040346-GAL4 | yak1694 | *D. yakuba* | attB/P integration |
| san2092_VT040346 | VT040346-GAL4 | san2092 | *D. santomea* | attB/P integration |
| yak1694_VT040347 | VT040347-GAL4 | yak1694 | *D. yakuba* | attB/P integration |
| san2150_VT040556 | VT040556-GAL4 | san2150 | *D. santomea* | attB/P integration |
| san2151_VT040556 | VT040556-GAL4 | san2151 | *D. santomea* | attB/P integration |
| yak2180_R71G01-AD | R71G01-p65ADZp | yak2180 | *D. yakuba* | attB/P integration |
| yak2177_R71G01-DBD | R71G01-ZpGAL4DBD | yak2177 | *D. yakuba* | attB/P integration |
| yak2177_R15A01-DBD | R15A01-ZpGAL4DBD | yak2177 | *D. yakuba* | attB/P integration |
| yak2177_R22D03-DBD | R22D03-ZpGAL4DBD | yak2177 | *D. yakuba* | attB/P integration |
| yak2177_VT040346-AD | VT040346-p65ADZp | yak2177 | *D. yakuba* | attB/P integration |
| yak2180_VT040346-DBD | VT040346-ZpGAL4DBD | yak2180 | *D. yakuba* | attB/P integration |
| yak2180_VT040347-AD | VT040347-p65ADZp | yak2180 | *D. yakuba* | attB/P integration |
| yak2177_VT040556-DBD | VT040556-ZpGAL4DBD | yak2177 | *D. yakuba* | attB/P integration |
| yakw; fru-GAL4 | yakfruGAL4, 3XP3::DsRed | UCSD 14021-0261.02 | *D. yakuba* | CRISPR knock-in |
| yakw; fru-AD | yakfrup65ADZp, 3XP3::DsRed | UCSD 14021-0261.02 | *D. yakuba* | CRISPR knock-in |
| yakw; fru-DBD | yakfruZpGAL4DBD, 3XP3::DsRed | UCSD 14021-0261.02 | *D. yakuba* | CRISPR knock-in |

**Supplementary Table 2:** Wild-type strains of the *D. melanogaster* species subgroup used in this study. The sample size of each strain used for characterizing song types was indicated.

| Species | Strain | Sample Size |
| --- | --- | --- |
| *D. melanogaster* | Oregon-R | 13 |
| *D. simulans* | sim5 | 14 |
| *D. mauritiana* | mau29 | 16 |
| *D. sechellia* | UCSD 14021-0248.07 | 19 |
| *D. yakuba* | UCSD 14021-0261.02 | 24 |
| *D. santomea* | UCSD 14021-0271.00 | 12 |
| *D. teissieri* | UCSD 14021-0257.01 | 20 |
| *D. orena* | UCSD 14021-0245.01 | 14 |
| *D. erecta* | UCSD 14021-0224.01 | 20 |

**Extended Data Figure 1:**
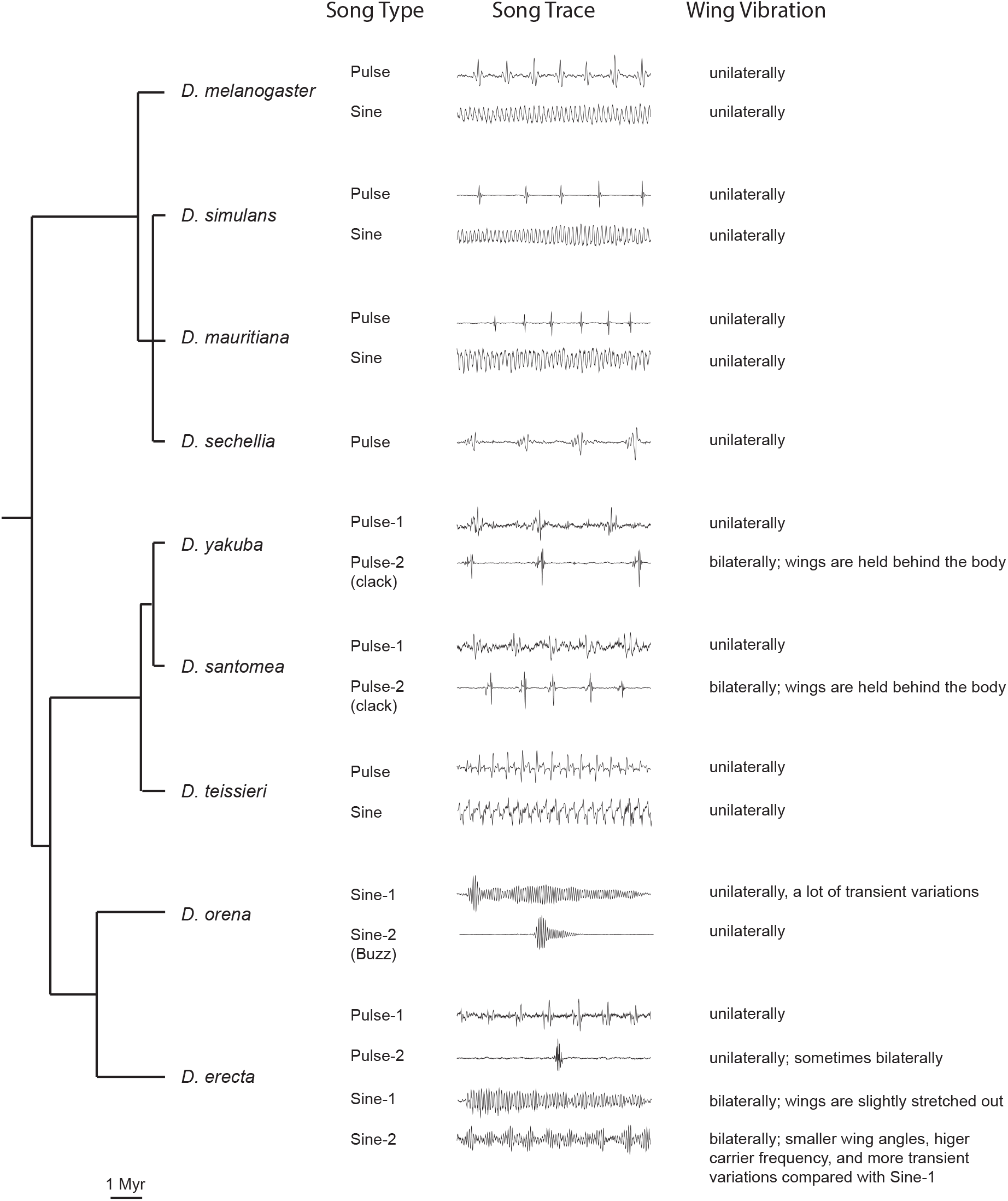
Summary of primary male courtship song types in the *D. melanogaster* species subgroup. For each type of courtship song, a 250 ms song trace is shown and the relevant wing motion is briefly described on the right. In general, pulse song is defined as song consisting of pulse events seperated by a relatively constant interval, and sine song is defined as continuous humming. Previous work reported that *D. orena* males produce pulse song consisting of polycyclic pulses^12^. We saw similar events ocassionally but the interval and the number of “pulse” cycles vary a lot within a song train, we therefore categorized these events into Sine-1, which exhibits a lot of transient variations. The inconsistency could be due to a difference either in definition or the particular strain used. *D. erecta* Pulse-2 events are not always organized into trains. SI Movie 1 illustrates each song type.

**Extended Data Figure 2:**
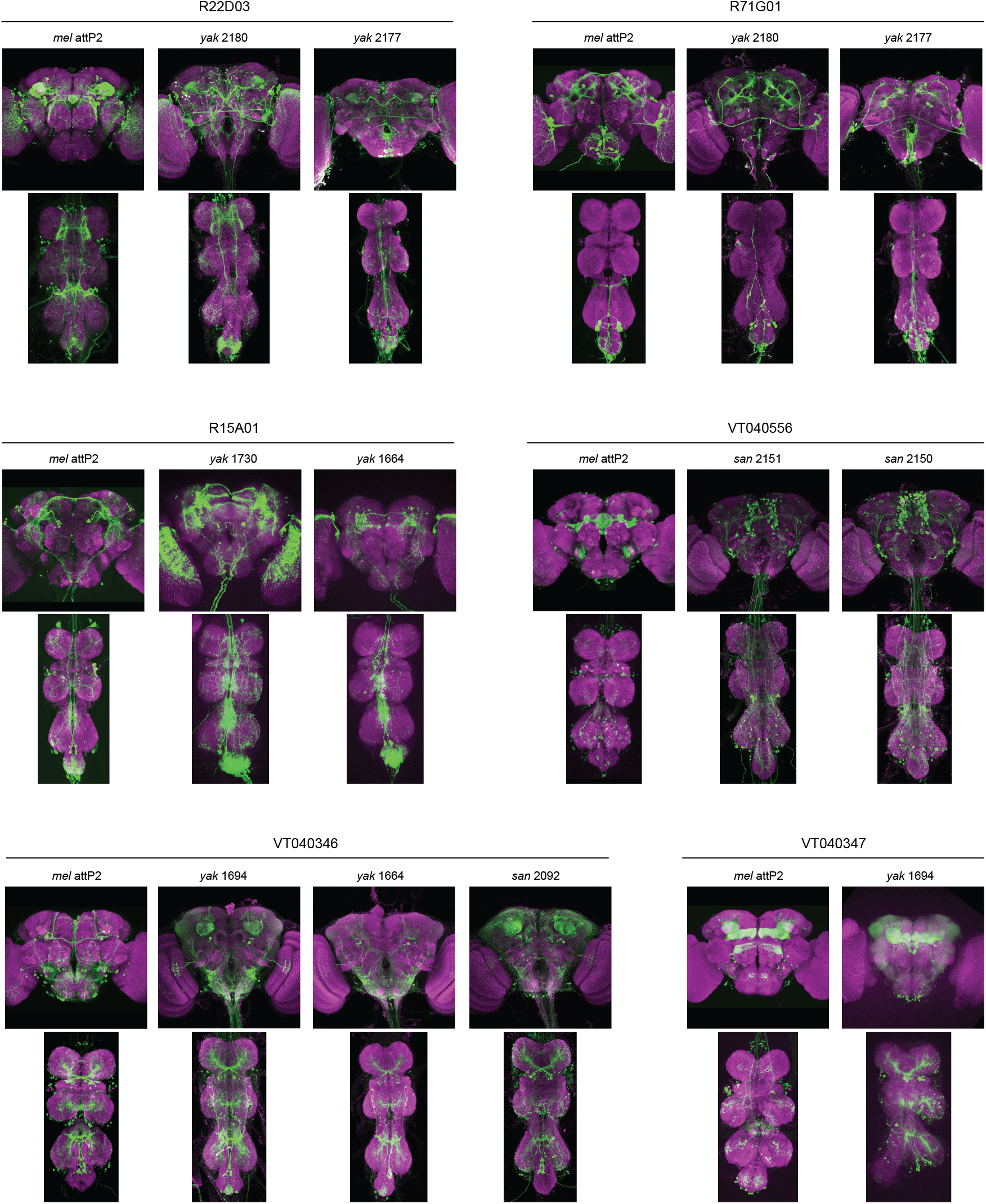
GAL4 reagents for P1 and plP10 neurons. Expression of GAL4 driven reporter gene (GFP or tdTomato) were visualized by staining the brains and VNSs of adult males using the corresponding antibody (green) and nc82 (magenta). The GAL4 ID, the species (*mel, D. melanogaster, yak, D. yakuba*; and *san, D. santomea*), and the attP site Into which It was stably Integrated are Indicated.

**Extended Data Figure 3:**
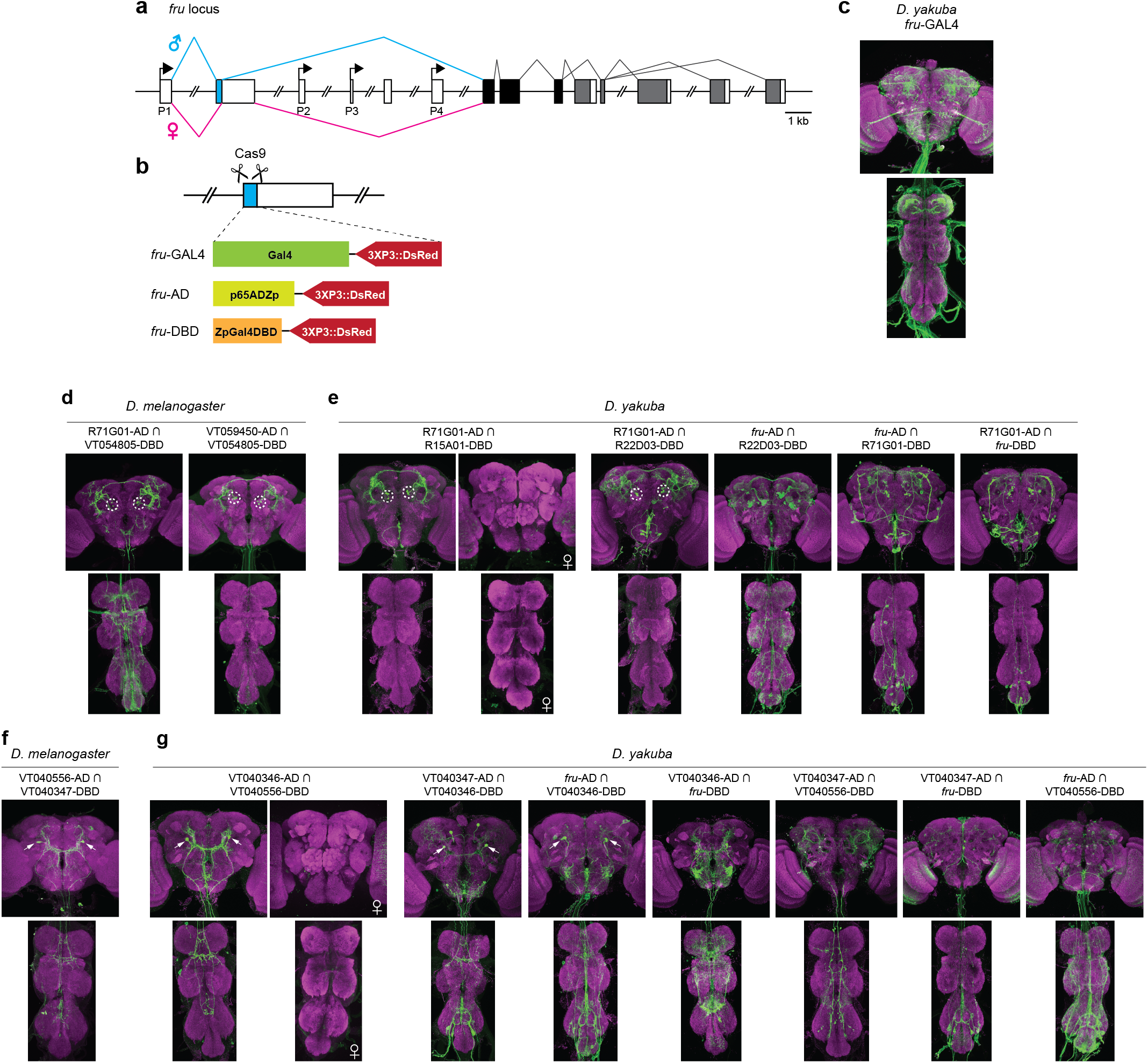
Split-GAL4 reagents of P1 and plP10 neurons. **a**, Schematic of *fru* gene structure. P1-P4 indicate alternative transcriptional start sites. Each box represents an exon. Open box Indicates untranslated region and filled box area Indicates protein coding region: cyan, male-specific coding exon; black: coding region of common exons; and grey: coding region of alternatively spliced exons, **b**, Generation of fru knock-¡n reagents via CRISPR-med¡ated HDR In *D. yakuba*. The male-specific exon (cyan) was replaced by GAL4, AD, and DBD, respectively, **c-g**, Immunosta¡n¡ng of brains and VNSs of adult flies for *D. yakuba* fru-GAL4 (**c**), P1 spllt-GAL4 lines (**d, e**), and plP10 spllt-GAL4 lines (**f, g**). Male brains and VNSs are shown unless Indicated. Expression of spllt-GAL4 driven reporter gene (GFP or tdTomato) were visualized by staining using the corresponding antibody (green) and nc82 (magenta). For spllt-GAL4 lines used In behavioral analysis, the cell bodies of the labelled neurons are marked with a dashed circle for P1 and an arrow for plP10.

**Extended Data Figure 4:**
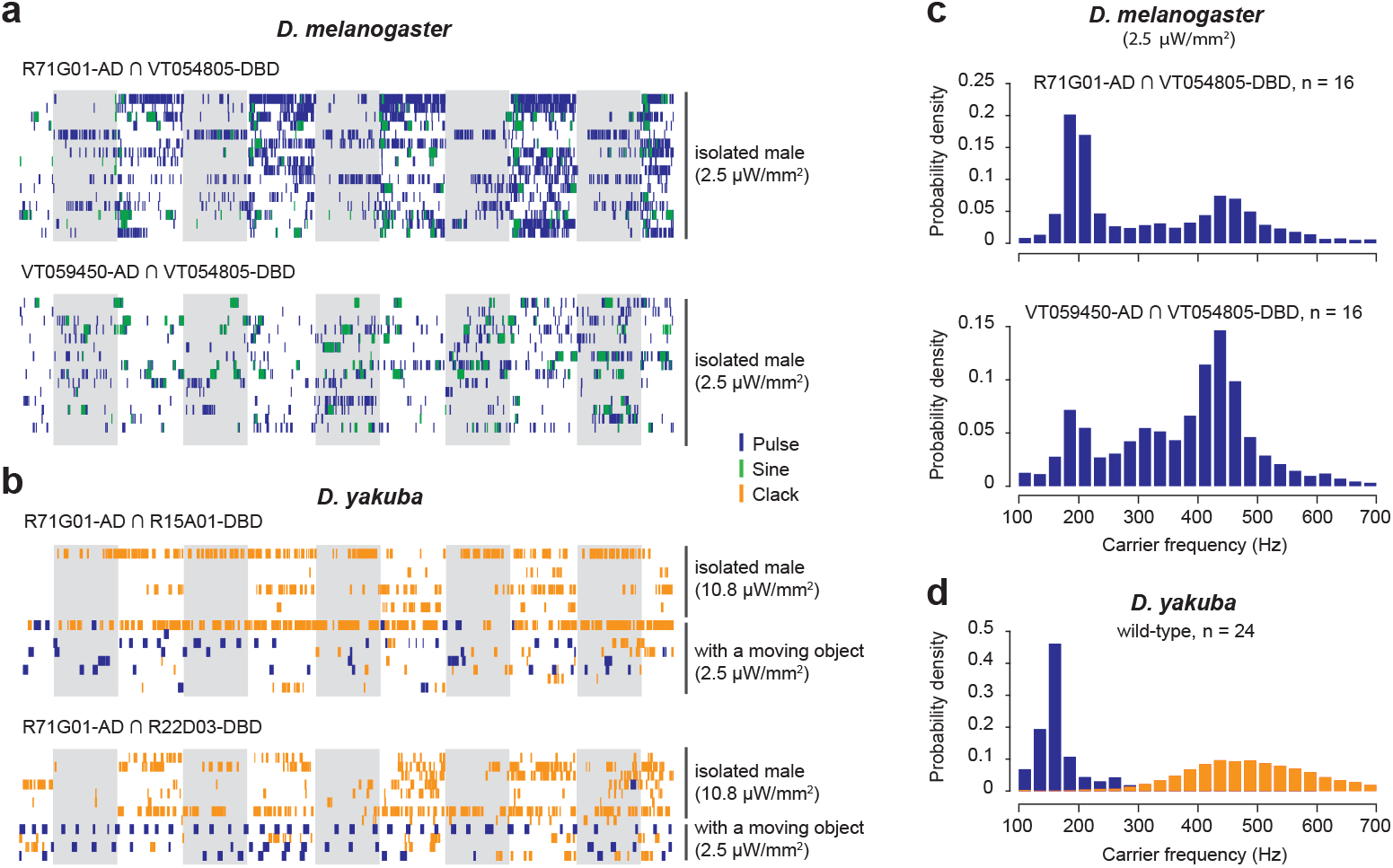
Song phenotypes resulting from P1 CsChrimson activation. **a, b**, Raster plot of song events induced by 635 nm red light illumination of *D. melanogaster* (**a**) and *D. yakuba* (**b**) males expressing CsChrimson in P1 neurons. Blue: pulse song; green: sine song; and orange: clack song. Each row represents song of one fly. Each grey bar represents a 30 s light stimulation trial. The trials shown here are part of an activation experiment using ramping irradiations and the trials at the irradiance level that gives relatively robust activation are shown, so song events that occur before the first stimulation shown here were induced by earlier stimulations that are not shown. *D. melanogaster* songs were analyzed with FlySong-Segmenter to identify pulse and sine events. *D. yakuba* songs were annotated manually to identify pulse, clack, and sine events, as the large amount of wing scissoring and abdomen quivering behaviors elicited by CsChrimson activation generated sound that confounded automatic song annotation. In *D. yakuba*, activation was performed on isolated males as well as in the presence of a moving object (a male). Repeating these experiments without red light illumination generated no (isolated male) or very little (with a moving object) song (data not shown). The activation phenotype of R71G01-AD ∩ R15A01-DBD is less robust than R71G01-AD ∩ R22D03-DBD, presumably because it labels only a small subset of P1 neurons (n=3 per hemisphere). **c**, The distribution of pulse carrier frequency in P1-activated *D. melanogaster* males using two split-GAL4 lines. **d**, The distribution of carrier frequency for pulse (blue) and clack (orange) song in wild-type *D. yakuba* males. Number of scored animals are indicated.

**Extended Data Figure 5:**
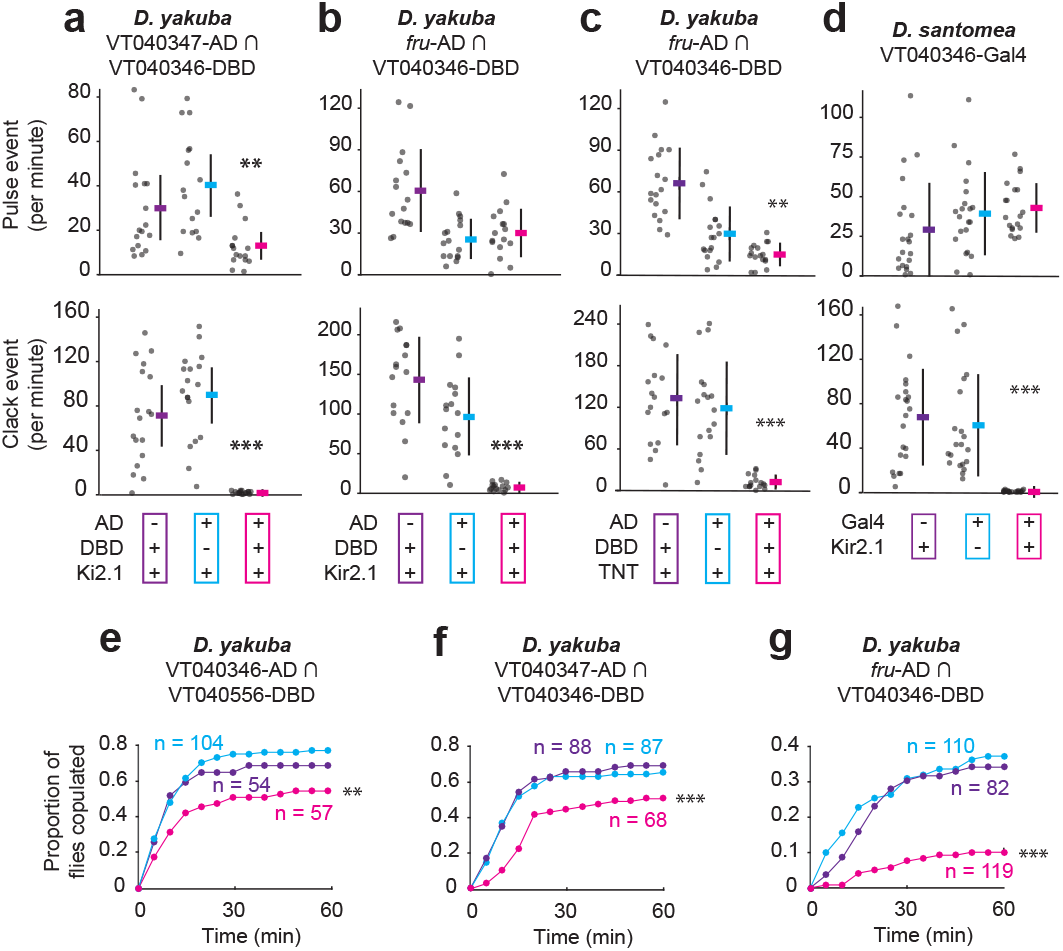
Song phenotypes and copulation rates resulting from pIP10 inactivation. **a-d**, Pulse and clack song production of pIP10 silenced males in *D. yakuba* using different split-GAL4 drivers and effectors (**a-c**) and in *D. santomea* using a non-sparse GAL4 driver (**d**). Data for each animal and mean ± SD are shown. n > 16. P values estimated with one-way ANOVA using a permutation test. **e-g**, Copulation success of pIP10 silenced males (pIP10 > TNT) in *D. yakuba*. The genotypes and their color representation are the same as in Fig. 2b and the panel a and b: AD control (purple), DBD control (cyan), and pIP10 silenced males (magenta). Sample size for each genotype is shown. *P* values measured via a logrank test. Significance is indicated only when the experiment group is significantly different from both control groups and the less significant result is shown. ***, *p* < 0.001; **, *p* < 0.01.

**Extended Data Figure 6:**
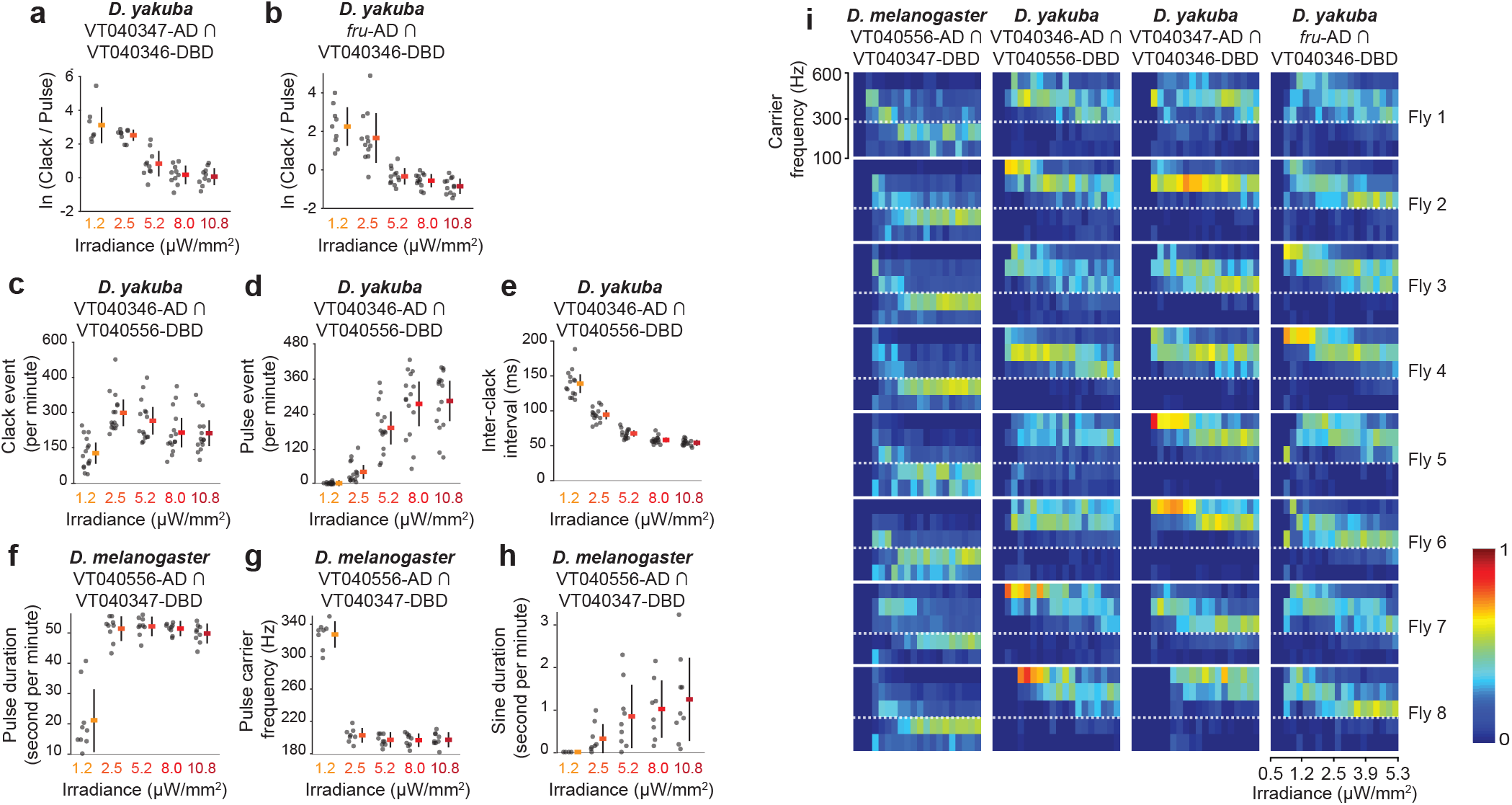
Additional song phenotypes resulting from pIP10 CsChrimson activation at different irradiances in *D. yakuba* and *D. melanogaster*. **a, b**, Natural log (ln) ratio of clack versus pulse events of pIP10 activated males in *D. yakuba* using the split-GAL4 lines VT040347-AD ∩ VT040346-DBD (**a**) and *fru-AD* ∩ VT040346-DBD (**b**). **c-e**, Song phenotypes of pIP10 activated males in *D. yakuba* using the cleanest pIP10 split-GAL4 driver (VT040346-AD ∩ VT040556-DBD): number of clack events per minute of activation (**c**), number of pulse events per minute of activation (**d**), inter-clack interval (**e**). **f-h**, Song phenotypes of pIP10 activated males in *D. melanogaster:* pulse duration per minute of activation (**f**), pulse carrier frequency (**g**), and sine duration per minute of activation (**h**). Data for each animal tested and mean ± SD are shown. n > 8. **i**, Heat map showing the distribution of song carrier frequency (pulse song for *D. melanogaster;* pulse and clack song combined for *D. yakuba*) of CsChrimson expressing pIP10 males with ramping irradiance using incremental steps of ~0.25 μW/mm^2^. Color represents the relative density of song events within a given carrier frequency range at the tested irradiance. See Fig. 2. for the mean of all tested animals.

**Extended Data Figure 7:**
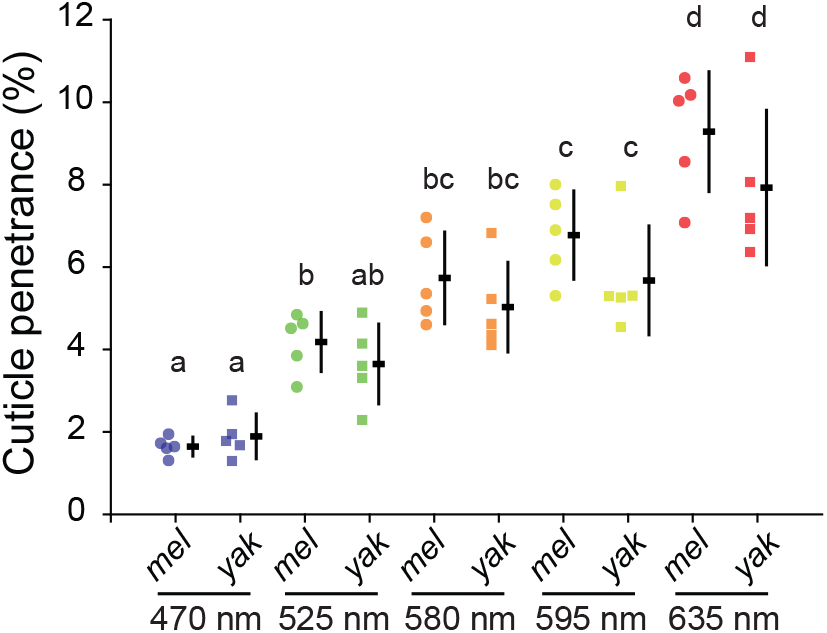
Penetrance of light through the cuticle in *D. melanogaster (mel)* and *D. yakuba (yak)* at different wavelengths. Data are shown for each animal and mean ± SD are plotted. n=5. Significance (*p* < 0.05) tested within and across species at each wavelength via two-way ANOVA and post-hoc Holm-Sidak pairwise multiple comparisons test. Same letter denotes no significant difference.

**Extended Data Figure 8:**
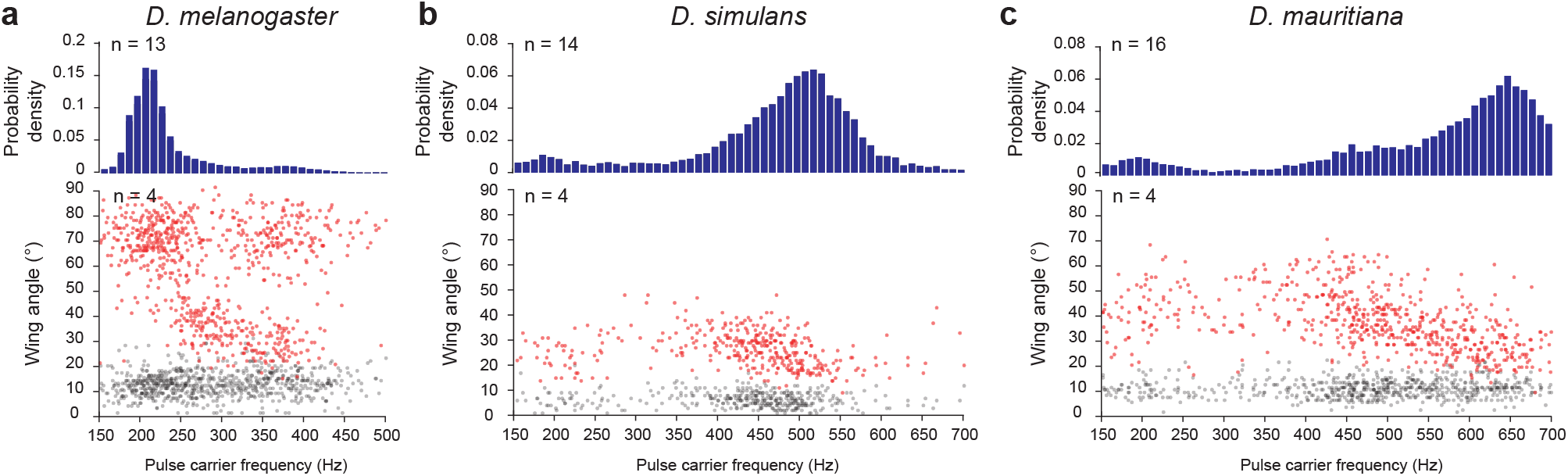
Carrier frequency and wing angle of pulse song in *D. melanogaster, D. simulans*, and *D. mauritiana*. **a-c**, Histograms of pulse carrier frequency from random samples of song (top plot) and wing angle versus pulse carrier frequency for selected songs (bottom plot) for *D. melanogaster* (**a**), *D. simulans* (**b**), and *D. mauritiana* (**c**). In the bottom plots, we oversampled pulses with less distributed carrier frequencies (see methods) to more fully characterize the assocation between wing angle and pulse carrier frequency. The angles of both wings were measured for each pulse event and the red and grey points show the angles of the more and less extended wings, respectively. Number of scored animals are indicated.

